# GenPerturb: sequence-grounded interpretation of perturbation transcriptomes using pretrained genomic models

**DOI:** 10.64898/2026.07.01.735806

**Authors:** Takuya Shihashi, Itoshi Nikaido

## Abstract

**Background:** Perturb-seq captures transcriptional responses to thousands of genetic and chemical perturbations, but does not directly resolve the cis-regulatory elements or transcription factor motifs underlying those responses. Existing approaches rely on indirect post hoc analyses or external epigenomic annotations, making it difficult to connect gene-level responses to specific regulatory element

**Results:** We present GenPerturb, a framework that leverages pretrained sequence-to-expression models to link perturbation-induced expression changes to candidate cis-regulatory elements. By contrasting perturbation and control states, GenPerturb prioritizes regulatory regions and transcription factor motifs associated with each perturbation. The model recapitulates perturbation-dependent gene expression patterns and enables sequence-level interpretation without requiring matched chromatin data. Across multiple perturbation types, GenPerturb identifies biologically meaningful regulatory programs, including lineage-specific and signaling-associated motif activities, even when corresponding transcription factor expression changes are limited.

**Conclusions:** GenPerturb converts gene-level expression responses from Perturb-seq into perturbation-specific, sequence-grounded cis-regulatory hypotheses. By prioritizing candidate regulatory elements and transcription factor motifs responsive to each perturbation without requiring matched chromatin data, GenPerturb enables mechanistic interpretation of transcriptional regulation and guides downstream experimental validation.

## Background

Large-scale perturbation profiling approaches such as Perturb-seq have enabled the measurement of transcriptional responses to hundreds to thousands of genetic perturbations at single-cell resolution [1–2]. However, Perturb-seq captures only the gene expression changes following perturbation and does not provide direct access to the underlying transcriptional regulatory states, such as which transcription factors are activated or which cis-regulatory elements are involved. In other words, modeling transcriptional regulatory states based solely on gene expression data is inherently limited, and additional information, including chromatin states and cis-regulatory context, is fundamentally required.

To address this limitation, a variety of approaches based on existing genome annotations and prior knowledge networks—such as motif enrichment analysis, enhancer–gene linking, and regulon inference—have been widely used [3–8]. While these methods provide effective strategies for indirectly inferring regulatory states, they often rely on annotations derived from specific cell types or steady-state conditions. As a result, they may not fully capture context-dependent regulatory states in newly generated datasets, particularly under perturbation conditions. Given these limitations, an ideal approach would be to simultaneously measure gene expression and chromatin states within the same cell, thereby directly linking transcriptional responses to their regulatory basis. As an experimental solution to this problem, Multiome Perturb-seq enables the simultaneous measurement of gene expression and chromatin accessibility in the same cell following perturbation [9–12]. This approach allows transcriptional responses to be directly associated with chromatin changes at regulatory regions. However, the addition of epigenomic measurements introduces constraints in terms of experimental cost and throughput, and cannot be applied retrospectively to the large body of existing RNA-only Perturb-seq datasets. Therefore, if cis-regulatory information could be computationally extracted from RNA-only perturbation data, it would be possible to extend analyses to a broader perturbation space, including existing datasets, without requiring additional experiments. Achieving this requires explicitly modeling the relationship between gene expression and the underlying sequence information.

A promising computational framework for this purpose is sequence-to-expression modeling, which directly predicts gene expression and chromatin states from DNA sequences. Models such as Enformer, Borzoi, and AlphaGenome leverage long-range sequence context spanning hundreds of kilobases and are capable of capturing regulatory sequence features associated with molecular phenotypes [13–15]. Through attribution analysis, these models can identify sequence elements that contribute to their predictions [16–17]. However, these models are primarily trained on steady-state data, and their attribution outputs reflect sequence features important for gene regulation under non-perturbed conditions.

Consequently, identifying regulatory elements that are newly engaged or repressed in response to perturbations requires a framework that explicitly extracts perturbation-dependent sequence contributions by comparing with control conditions.

To address this challenge, we propose GenPerturb, an analytical framework for deriving sequence-grounded regulatory hypotheses from observed Perturb-seq responses (Fig. 1). GenPerturb uses pretrained sequence-to-expression models as regulatory sequence representations and adapts them to a target RNA-only Perturb-seq dataset by fitting observed expression outputs for the control condition and each perturbation condition. For each gene, control and perturbation outputs are fitted from the same local genomic sequence. This places fitted expression responses and attribution profiles on a shared gene locus and sequence coordinate system, enabling the identification of candidate regulatory elements and motif instances.

**Figure 1.**
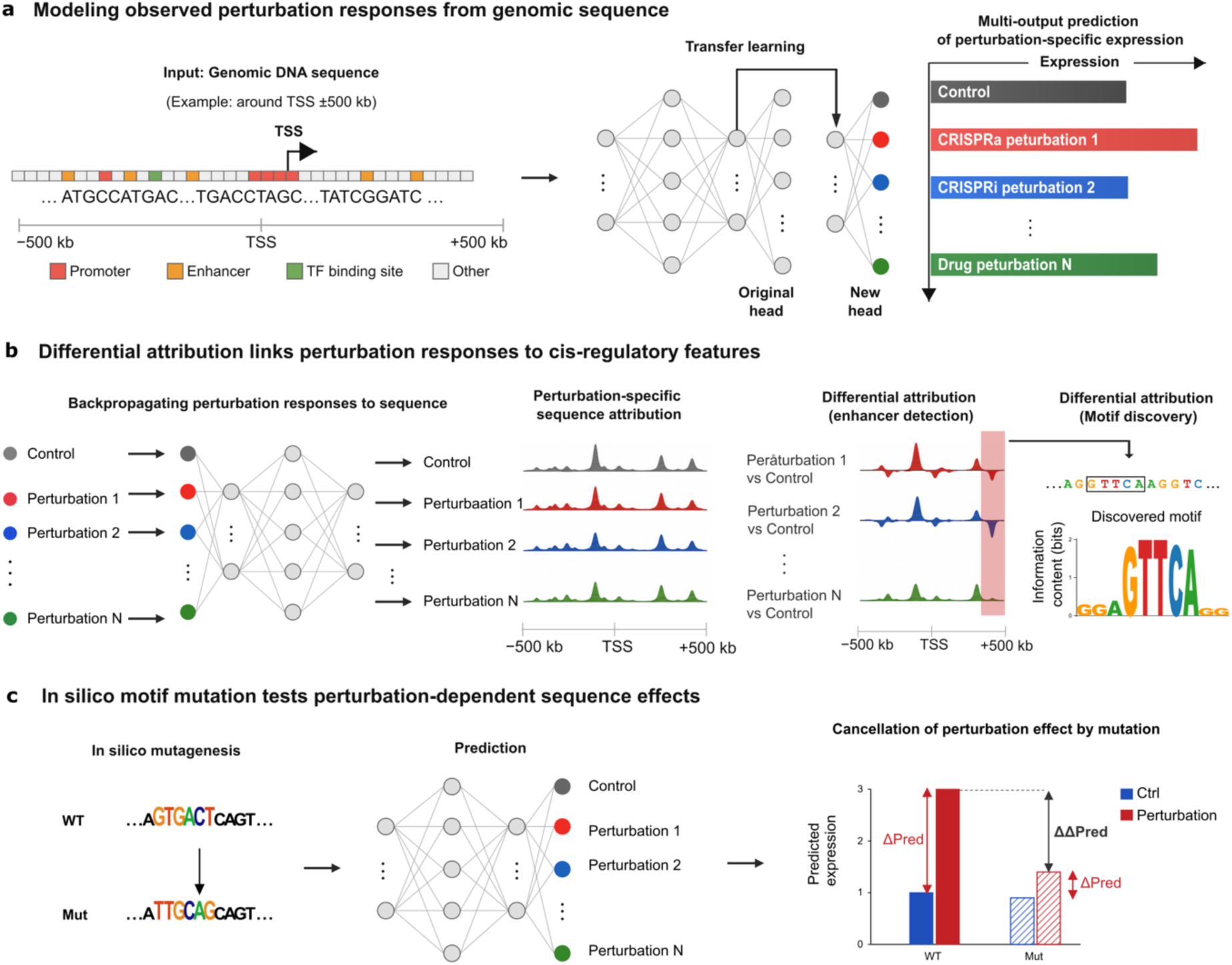
GenPerturb links observed perturbation responses to cis-regulatory sequence features. a, Overview of transfer learning-based sequence modeling of observed perturbation responses. Genomic DNA sequence around each gene, illustrated here as a window centered on the transcription start site, is used as input to GenPerturb. Pretrained sequence representations are transferred to the target Perturb-seq dataset by fitting condition-specific output channels corresponding to control and observed perturbation states, adapting regulatory sequence features to the target assay, cell state, and expression scale while representing all outputs on a shared gene locus and sequence coordinate system. b, Differential attribution for nominating and ranking perturbation-associated cis-regulatory candidates. Output-to-input attribution is computed separately for control and perturbation-specific outputs by backpropagating each condition output to the input sequence. Subtracting the control attribution profile from each perturbation attribution profile yields differential attribution tracks, which are used to rank candidate regulatory regions and motif-level sequence patterns associated with fitted perturbation responses. c, In silico motif mutation for evaluating model-internal sequence sensitivity. Ranked motif instances are mutated in the input sequence, and changes in fitted control and perturbation outputs are compared between wild-type and mutated sequences. Reduction of the perturbation-associated output difference after motif mutation supports a model-internal contribution of the candidate sequence feature to the fitted perturbation response.

GenPerturb then computes differential attribution scores by subtracting control attribution profiles from perturbation attribution profiles, producing perturbation-minus-control attribution tracks [18–19] that highlight sequence positions associated with the fitted perturbation response. These tracks are summarized as ranked candidate regulatory elements and motif instances for each gene–perturbation pair. In this way, GenPerturb turns RNA-only Perturb-seq responses into locus-resolved, sequence-grounded regulatory hypotheses without requiring matched ATAC-seq, ChIP-seq, or multiome data from the target experiment (see Supplementary Table 1 for the positioning of our method relative to existing approaches).

Here, we demonstrate the ability of GenPerturb to extract sequence-grounded regulatory information from Perturb-seq datasets without requiring matched chromatin measurements in the target experiment. First, we show that the fitted sequence-based model retains perturbation-dependent expression structure across multiple Perturb-seq datasets. Comparison with perturbation multiome data further demonstrates that differential attribution can rank candidate cis-regulatory regions and transcription factor motifs with support comparable to, and in aggregate modestly exceeding, annotation-based methods.

Application of GenPerturb to regulator perturbations along the erythroid and granulocyte differentiation lineages reveals that fitted outputs retain lineage-associated structure and that motif-level attribution captures activity-like regulatory signals along these differentiation axes. Finally, analysis of compound perturbations shows that motif-level attribution can recover activity-like regulatory programs linked to ligand-dependent receptor activation, stress-response signaling, and inflammatory signaling, even when matched or motif-associated transcription factor mRNA levels change only weakly, supporting regulatory interpretation of compound effects through ranked candidate programs.

## Results

### GenPerturb anchors control and perturbation responses to shared sequence coordinates for differential attribution

GenPerturb was designed to model perturbation-resolved expression responses within a shared genomic sequence framework (Fig. 1). For each gene, pretrained sequence representations are fitted to control and perturbation-resolved expression outputs, so that all condition-specific outputs can be compared on the same genomic input (Fig. 1a; Fig. S1). After fitting, GenPerturb compares output-to-input attribution profiles between perturbation and control outputs on the same sequence coordinates. This control-subtracted differential attribution provides the basis for ranking perturbation-associated candidate regions and motif-level sequence patterns (Fig. 1b).

We further use in silico motif mutation as a model-internal sensitivity analysis, testing whether ranked motif instances alter the fitted perturbation-control difference (Fig. 1c). For each gene-perturbation pair, GenPerturb thus produces ranked candidate enhancer-like regions and motif instances that can be used for downstream regulatory interpretation and integration with existing enhancer or motif annotations.

To support this framework, we evaluated GenPerturb in three complementary validation layers throughout the remainder of the study: (i) retention of perturbation-dependent fitted expression structure across diverse Perturb-seq datasets (Fig. 2); (ii) agreement of differential attribution with independent multiome-derived chromatin and motif evidence, benchmarked against rE2G extended, rE2G, ABC, and TSS-distance baselines (Fig. 3); and (iii) sensitivity of fitted perturbation responses to counterfactual motif disruption by in silico mutation of ranked motif instances (Fig. 3). Together these layers test whether expression structure, sequence-level priors, and sequence-level counterfactuals each support the ranked candidate regulatory features. Lineage-associated and compound perturbation analyses (Figs. 4, 5) then illustrate how the resulting motif-level signals can capture activity-like regulatory variation that is not fully explained by transcription factor mRNA abundance.

**Figure 2.**
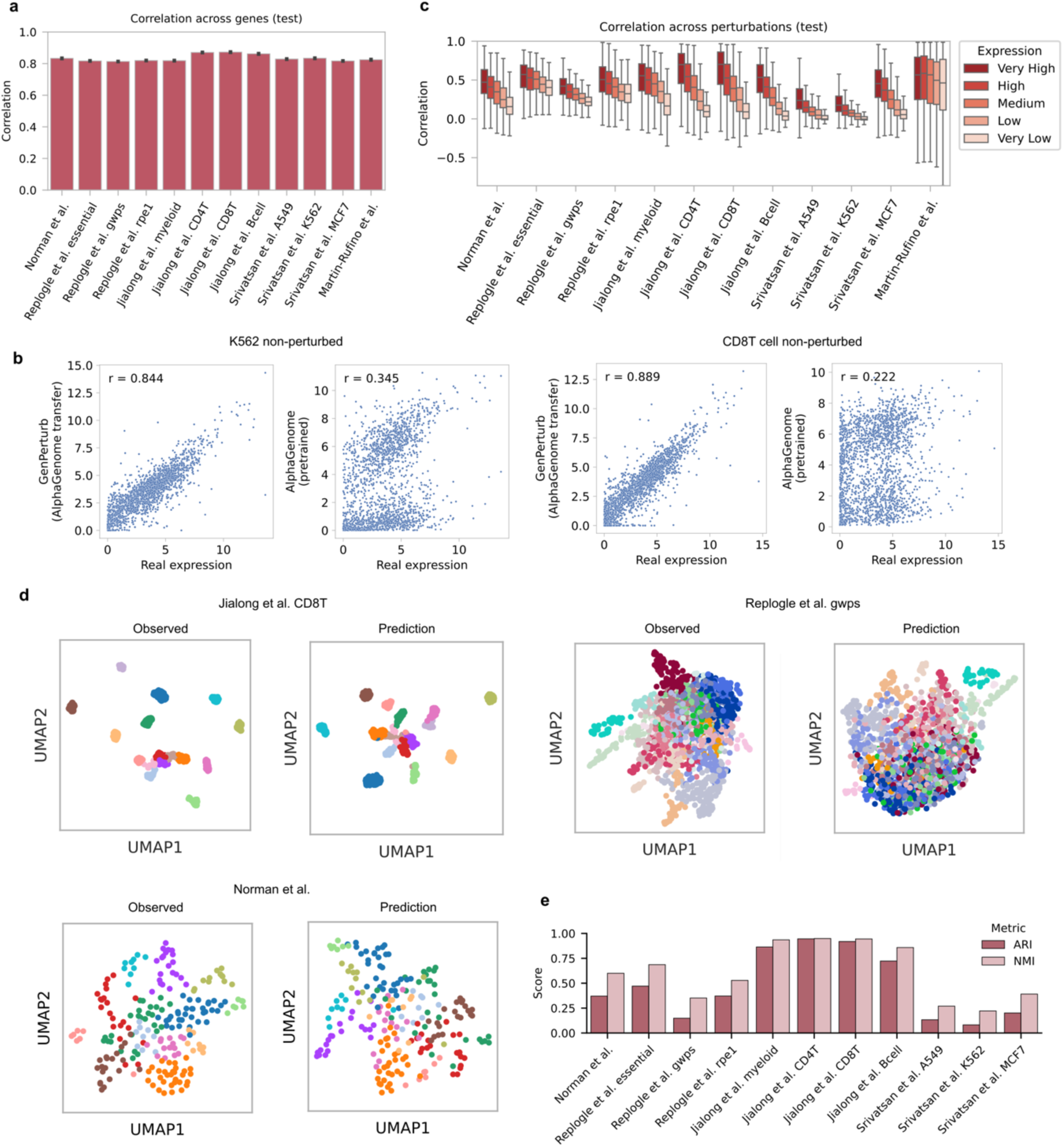
GenPerturb retains perturbation-dependent expression structure after fitting to observed Perturb-seq data. a, Across-gene agreement between observed expression and fitted GenPerturb outputs for each perturbation condition across Perturb-seq datasets (Pearson r across genes, computed per perturbation condition; held-out test split). Bars show correlation across genes within each condition, summarized for each dataset. b, Comparison of observed control expression with fitted outputs from the transferred AlphaGenome-based GenPerturb model and baseline non-perturbed outputs from the original pretrained AlphaGenome prediction head, in K562 and CD8 T cells (Pearson r across genes). c, Across-perturbation agreement for individual genes (Pearson r across perturbations, computed per gene; expression quintiles; held-out test split). For each gene, observed and fitted expression values were compared across perturbation conditions, showing stronger agreement for highly expressed genes (roughly r = 0.3-0.7 across datasets) and weaker agreement for lowly expressed genes (roughly r = 0.1-0.5). d, Low-dimensional organization of observed and fitted expression profiles for datasets with pronounced perturbation structure in the observed space. Dots in both observed and fitted UMAPs are colored by Leiden clusters computed from the observed expression space; clustering of the same colors in the fitted UMAP indicates preservation of observed perturbation-state structure. e, Cluster concordance between observed and fitted expression spaces across datasets (ARI, NMI; observed vs fitted Leiden partitions; held-out test split).

**Figure 3.**
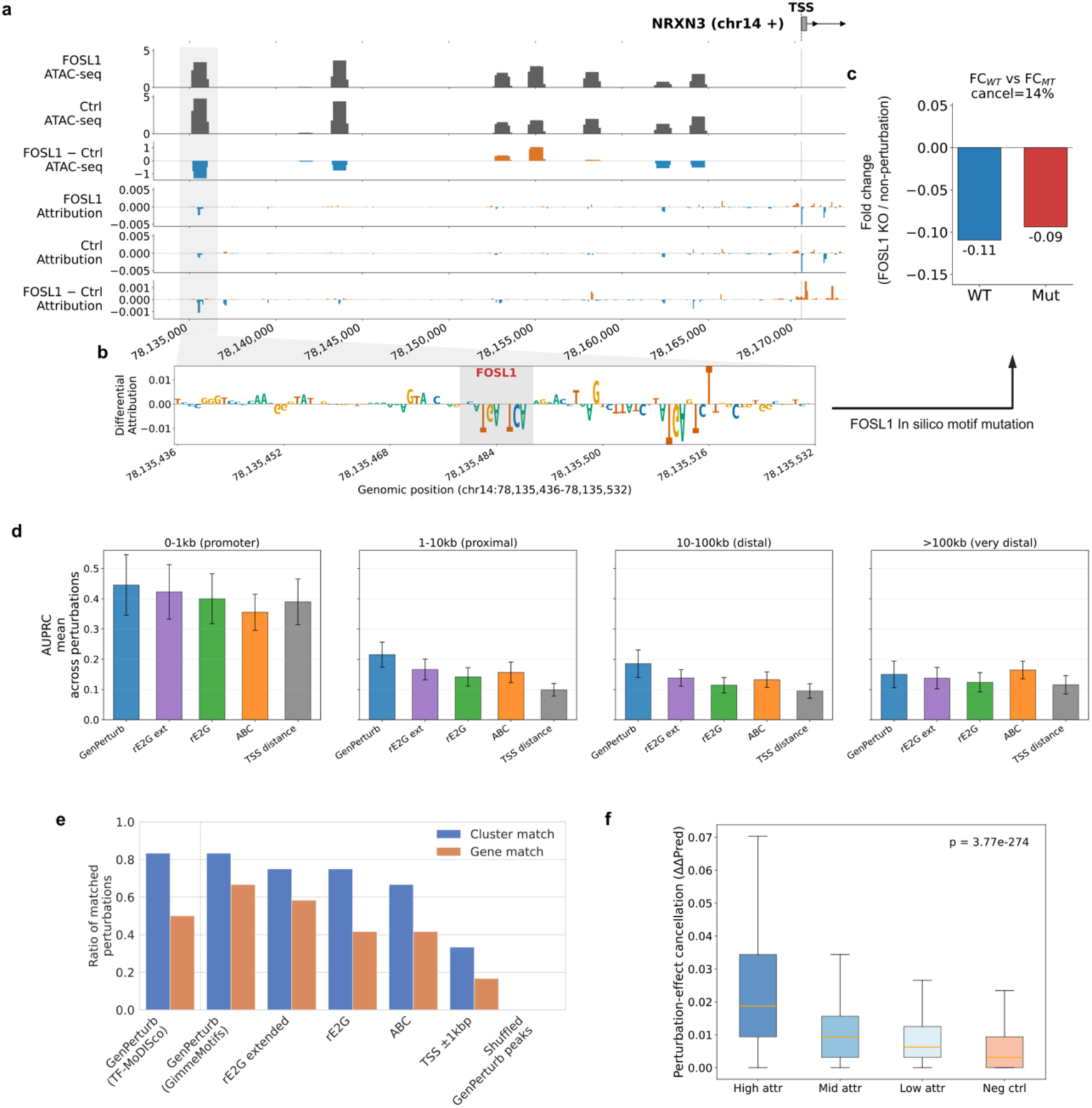
Differential attribution prioritizes perturbation-associated cis-regulatory elements and motifs. a, Representative locus-level example showing perturbation-resolved ATAC-seq signal, condition-specific attribution, and control-subtracted differential attribution at the NRXN3 locus under FOSL1 perturbation. Differential attribution highlights discrete noncoding regions within the same genomic coordinate system used for control and perturbation outputs. b, Sequence-level zoom of a prioritized region, showing differential attribution over nucleotide positions and the corresponding motif instance (FOSL1). c, Model-internal in silico motif mutation for the representative locus. Randomizing the prioritized motif instance partially attenuates the fitted perturbation-associated expression change, using mutant predictions averaged across five mononucleotide-preserving random shuffle seeds. d, Enhancer prioritization benchmark using perturbation-resolved TF-sensitive accessible chromatin regions (Martin et al. Table S2) as reference positives (AUPRC; mean across perturbations; |log2FC| ≥ 0.5 for gene expression fold change; ≥10 positive peaks per distance-based bin). Differential attribution, rE2G extended, rE2G, ABC, and TSS-distance scores were compared across promoter-proximal (<1 kb), proximal (1–10 kb), distal (10–100 kb), and very distal (>100 kb) regulatory regions. Bars show the mean AUPRC across perturbations and whiskers the across-perturbation standard error of the mean (SEM), with each perturbation contributing one observation per stratum. Of the 12 Martin et al. perturbation conditions, only those with at least 10 positive peaks within a given distance stratum were evaluated, yielding 7 perturbations per stratum and 28 perturbation × distance strata in total. Performance varied across perturbations and distance strata. e, Motif recovery analysis comparing attribution-derived motifs and annotation-based regulatory regions (evaluated separately for each perturbation and motif-discovery method; TF-MoDISco and GimmeMotifs enrichments filtered at q ≤ 0.05; gene match or JASPAR transcription factor family cluster match to the perturbed transcription factor). Two complementary procedures were applied to GenPerturb attribution: (i) TF-MoDISco directly on differential attribution profiles to obtain attribution-derived seqlet motifs, and (ii) GimmeMotifs known-motif enrichment on attribution peak regions within the same unified motif-enrichment framework applied to annotation-based regions (rE2G extended, rE2G, ABC, TSS-proximal). Bars indicate the fraction of perturbations for which motifs corresponding to the perturbed transcription factor were successfully recovered. f, In silico mutation analysis of TF-MoDISco-identified motif instances whose Tomtom annotation matches the perturbed transcription factor or its JASPAR cluster (q ≤ 0.05), stratified by differential attribution strength (5 mononucleotide-preserving random shuffle seeds averaged per mutated interval; groups = tertiles of attribution strength plus paired length-matched negative controls at motif-absent loci; one-sided Mann-Whitney U for high vs negative control where shown).

**Figure 4.**
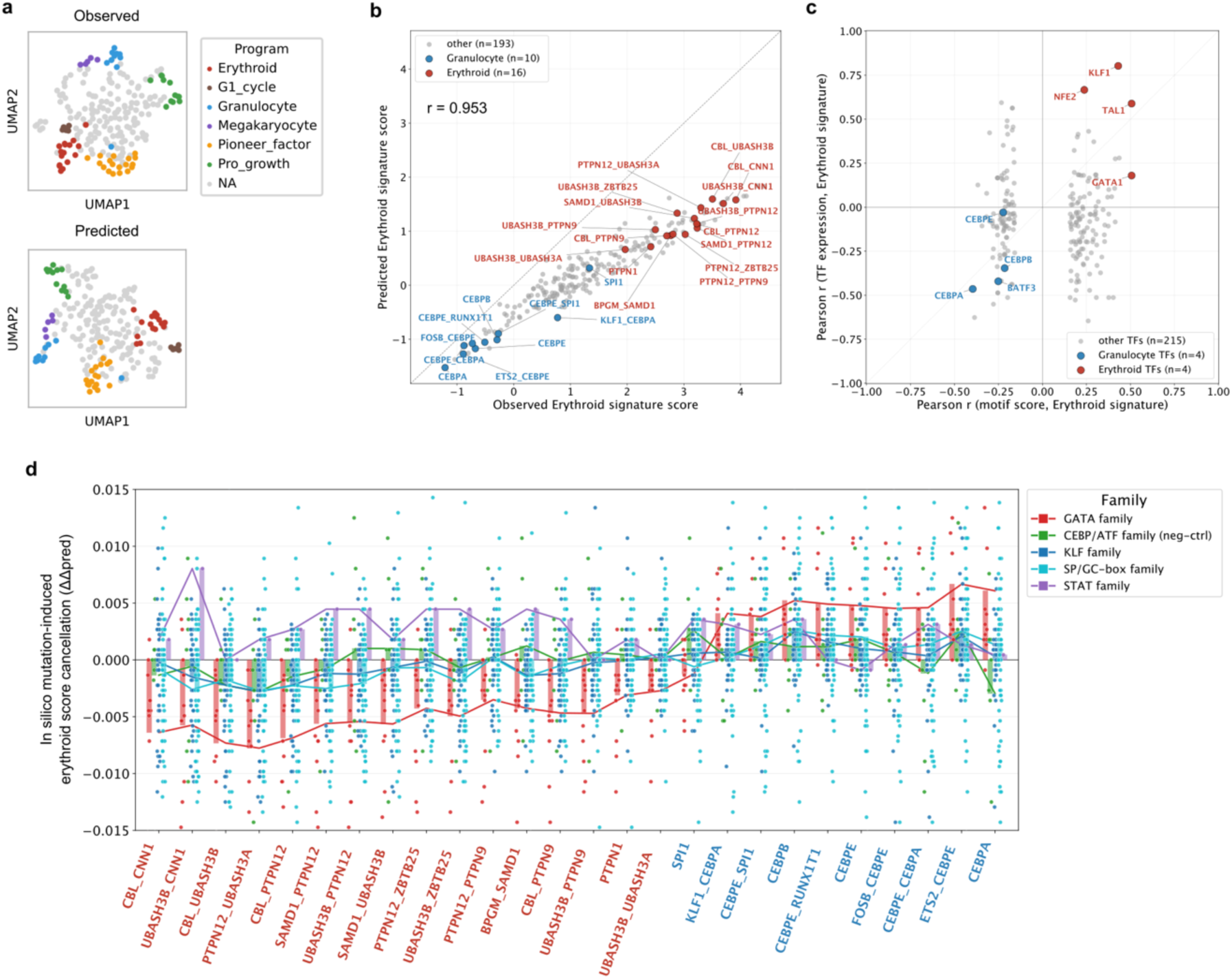
GenPerturb links differentiation-associated expression axes to motif-level regulatory associations. a, Low-dimensional organization of observed and fitted expression profiles annotated by lineage-related perturbation programs from the original Norman study. Each dot represents a perturbation and is colored by the original study annotation. Structured lineage-associated organization observed in the original data is retained in the fitted expression space. b, Comparison of observed and fitted erythroid signature scores across perturbations using the erythroid signature from the original Norman study (Pearson r). Each dot represents a perturbation; red dots indicate perturbations annotated as erythroid differentiation and blue dots indicate perturbations annotated as granulocyte differentiation in the original study. c, Relationship between TF-MoDISco-derived motif significance scores (-log10 q) and TF mRNA expression along the erythroid axis. For each motif or TF family, the y-axis shows the correlation between TF mRNA expression and the erythroid signature score (Pearson r), and the x-axis shows the correlation between the motif significance score and the erythroid signature score. d, In silico mutation of erythroid marker gene–associated and TF-MoDISco–identified motif instances (5 mononucleotide-preserving random shuffle seeds averaged per motif instance). Each dot represents a motif instance, and the y-axis shows the change in erythroid marker-gene signature score after motif mutation (ΔΔpred). Larger negative values indicate stronger cancellation of perturbation-associated erythroid signature changes following motif disruption. X-axis labels are colored using the original Norman study annotations as in panel b, with erythroid differentiation perturbations in red and granulocyte differentiation perturbations in blue.

### GenPerturb captures perturbation-dependent expression structure across diverse Perturb-seq datasets

Before using differential attribution to rank perturbation-associated cis-regulatory candidates, we first asked whether the transferred sequence-based model retained the expression structure required for comparing control and perturbed conditions. Using AlphaGenome as the primary backbone [15], we applied GenPerturb to twelve Perturb-seq datasets from Norman et al., Replogle et al., Martin et al., Jiang et al., and Srivatsan et al., spanning CRISPR activation, CRISPR interference, CRISPR knockout, and compound perturbations across both genetic and chemical perturbation modalities [12,20–23] (Supplementary Table 2). As an initial assessment, fitted expression profiles showed high agreement with observed expression across genes within each perturbation condition, with correlations consistently around 0.8 or higher across datasets (Fig. 2a). This result indicates that dataset-specific transfer preserves the dominant gene-level expression structure of each perturbation state. Because across-gene evaluation is strongly influenced by stable basal expression differences among genes, we next asked whether transfer to the target Perturb-seq dataset was necessary. Under non-perturbed conditions, direct use of the original pretrained AlphaGenome output showed substantially lower agreement with observed expression than the transferred model. For example, across-gene correlation improved from r = 0.345 to r = 0.844 in K562 and from r = 0.222 to r = 0.889 in CD8 T cells after transfer (Fig. 2b). These results indicate that pretrained sequence representations cannot be used as static expression outputs for Perturb-seq interpretation, but instead must be adapted to the target assay, cell state, and expression scale.

We then evaluated whether the transferred model retained perturbation-dependent variation at the level of individual genes. In contrast to the across-gene analysis, this evaluation measures whether each gene shows concordant variation across perturbation conditions, and therefore more directly tests the condition-dependent structure required for downstream differential attribution. Agreement was strongest for highly expressed genes and in datasets with strong perturbation structure (roughly r = 0.3-0.7), whereas lowly expressed genes showed weaker correspondence (roughly r = 0.1-0.5; Fig. 2c). To determine whether this across-perturbation signal could be explained by trivial expression structure, we compared GenPerturb with simple baselines. Both the control-copy and perturbation-mean baselines showed high across-gene correlations, confirming that this metric alone can be dominated by stable expression differences among genes. However, these baselines failed to reproduce gene-wise variation across perturbations, with across-perturbation correlations compressed near zero for most datasets and expression bins (Fig. S2). This calibration against simple baselines follows recent work emphasizing that mean-like predictors can appear competitive under poorly calibrated perturbation-response metrics [24]. Thus, the signal observed in Fig. 2c is not explained by simply reusing the non-perturbed expression profile or by applying a shared average response across perturbations. This supports the use of condition-specific predicted outputs for downstream differential attribution.

We further examined whether the global organization of perturbation states was preserved in low-dimensional expression space. Observed and predicted profiles showed broadly concordant organization. Datasets with pronounced perturbation structure in the observed space, such as Jiang et al. CD8 T cells and Norman K562, retained similarly separated structure in the predicted space (Fig. 2d), whereas datasets with more diffuse observed structure, such as the Srivatsan compound perturbation datasets, showed correspondingly weaker separation (Fig. S3). Cluster concordance metrics showed consistent trends across datasets, supporting that GenPerturb retains not only gene-wise expression structure but also the global geometry of perturbation-induced transcriptional states (Fig. 2e).

Finally, we examined the contribution of pretrained sequence representations and the stability of the transfer procedure. In comparisons across Enformer [13], Borzoi [14], AlphaGenome [15], and a Simple CNN trained without large-scale regulatory pretraining, AlphaGenome-derived representations showed stronger agreement both in correlation-based evaluations and in low-dimensional perturbation geometry (Figs. S4, S5). Multifold evaluation using fold-specific pretrained AlphaGenome checkpoints and matched genomic splits showed consistent results across the tested settings, supporting the robustness of the retained perturbation-dependent structure to the evaluated split and checkpoint choices (Fig. S6). We also compared full fine-tuning, LoRA [25], and feature extraction, which yielded comparable agreement across evaluation metrics. We therefore used feature extraction in subsequent analyses to reduce computational cost while retaining the perturbation-dependent expression structure required for attribution analysis (Fig. S7). This expression-level evaluation provides the first layer of validation for GenPerturb: fitted expression profiles retain perturbation-dependent structure before sequence-level interpretation.

### Differential attribution prioritizes perturbation-associated cis-regulatory elements

Having established that GenPerturb preserves perturbation-dependent expression structure, we next examined whether differential attribution can highlight cis-regulatory elements associated with perturbation responses. This step is designed to go beyond post hoc motif enrichment of differentially expressed genes by asking which sequence elements and motif instances at the corresponding gene loci are candidates for contributing to each fitted perturbation-associated response. To define an external benchmark for enhancer-oriented validation, we surveyed publicly available perturbation multiome resources. Although several candidate datasets contained paired RNA and chromatin measurements, most could not be used for quantitative perturbation-resolved enhancer evaluation because perturbation identities, sufficient perturbation-associated expression changes, or corresponding perturbation-associated ATAC changes were not available at the required scale (Supplementary Table 3). We therefore used the Martin et al. erythroid Perturb-multiome dataset as the primary external benchmark, as it provides perturbation-resolved chromatin accessibility together with enhancer-gene reference information [12]. Although this restricts the benchmark to 12 perturbation conditions, it is the largest publicly available resource that satisfies our perturbation-resolved AUPRC evaluation criteria. This limited availability of usable perturbation-multiome benchmarks also illustrates the practical motivation for GenPerturb: in many RNA-only perturbation studies, direct chromatin measurements are unavailable, and sequence-based hypothesis prioritization can provide a regulatory interpretation layer for selecting candidate elements and motifs for follow-up analyses.

Before evaluating enhancer-level agreement, we first asked whether control-subtracted differential attribution provided an appropriate ranking signal for perturbation-associated responses. In the Martin dataset, differential attribution showed stronger enrichment for genes with large observed perturbation responses than either control-output attribution or perturbation-output attribution alone, particularly at stringent top-gene cutoffs. Across perturbations within each gene, differential attribution also showed higher concordance with observed absolute fold-change than perturbation-output attribution without control subtraction, especially among genes with larger response magnitudes (Fig. S8). This supports the use of control-subtracted attribution as the ranking signal for downstream regulatory prioritization, while leaving enhancer-level validation to the independent chromatin benchmark.

As an illustrative locus-level example before aggregate benchmarking, we examined NRXN3 under FOSL1 perturbation (Fig. 3a-c). Differential attribution highlighted a discrete noncoding region whose signal differed between control and perturbation outputs. This region overlapped a FOSL1-related motif instance identified by TF-MoDISco, consistent with the perturbed regulator analyzed at this locus. In silico mutation of this motif instance partially attenuated the model-derived perturbation-associated expression changes, illustrating how sequence features highlighted by differential attribution can be connected to fitted responses within the model (Fig. 3a-c).

Having established differential attribution as the ranking signal, we next asked whether the resulting candidate regions were supported by independent perturbation-resolved chromatin evidence. We compared differential attribution against rE2G extended, rE2G, ABC, and TSS-distance baselines using perturbation-resolved AUPRC evaluation [4–5]. Across the aggregate evaluation, GenPerturb achieved a small but consistent advantage over the strongest annotation-based comparators, and the advantage was clearer in distal regions where promoter-proximal motif enrichment and distance-based linking alone provide limited resolution (Fig. 3d). The differences are modest in absolute magnitude, and we interpret them as evidence that an RNA-only sequence-fitted model can match or modestly exceed annotation-heavy enhancer-linking baselines on perturbation-sensitive accessible chromatin, rather than as a claim of broadly superior enhancer prediction. ABC and rE2G provide strong annotation-based comparators when matched epigenomic resources are available, whereas GenPerturb does not require a precomputed enhancer-gene map for the target RNA-only perturbation experiment and instead derives candidate regulatory regions directly from the perturbation expression data itself. Per-perturbation analysis across promoter-proximal and distal strata showed that performance depended strongly on the number of evaluable positive regions per perturbation (Fig. S9). Differential attribution was often among the strongest scoring schemes when many positive enhancer candidates were available. In contrast, for perturbations with sparse positive regions, ABC or rE2G extended sometimes performed better, consistent with the increased sensitivity of AUPRC under sparse-positive settings and with the different information sources used by each method. These results argue against a single universally optimal enhancer score. Instead, they support a complementary use of model-derived differential attribution and annotation-based enhancer-linking approaches, with the most informative score depending on the perturbation, genomic distance stratum, and density of evaluable regulatory changes (Fig. 3d; Fig. S9).

We next assessed motif recovery using two complementary procedures. First, TF-MoDISco was applied directly to differential attribution profiles to extract recurring sequence patterns from attribution-derived regions [26], with motif identities assigned using Tomtom from the MEME Suite [7–8]. Second, to enable a like-for-like comparison with ABC and rE2G, we performed GimmeMotifs known-motif enrichment on each candidate-CRE region set under a unified framework, applying the same enrichment procedure to GenPerturb-derived regions and annotation-based regions [3–5]. The TSS ± 1 kb analysis represents the closest promoter-proximal counterpart to conventional post hoc motif enrichment over responsive genes, whereas ABC and rE2G extend the same enrichment framework to annotation-based enhancer-linked regions. Under this unified GimmeMotifs framework, GenPerturb-derived regions yielded a small but consistent advantage over the annotation-based region sets in recovering motifs corresponding to the perturbed transcription factor or its JASPAR cluster (cluster-matched ratio 0.83, 10/12 perturbations, for GenPerturb-derived regions vs 0.75, 9/12, for the next-best annotation-based region set, rE2G extended), and TF-MoDISco directly on attribution recovered overlapping motif identities (cluster match 0.83, 10/12) [27]. Similar results were reproduced in an independent CRISPRa dataset, with consistently strong recovery performance observed across both CRISPR KO and CRISPRa perturbations (Fig. 3e; Fig. S10a).

To further examine the functional relevance of these motifs within the model, we performed in silico motif mutation. For each motif instance, the sequence was randomized and the resulting change in model output was measured. Stronger differential attribution was associated with larger output changes upon mutation. In contrast, regions lacking motif instances or showing low attribution strength exhibited little to no such relationship when used as negative controls. These results support a model-internal correspondence between attribution magnitude and sequence sensitivity, rather than serving as direct experimental evidence of enhancer function (Fig. 3f; Fig. S10b). Together with the expression-structure analyses in Fig. 2 and the independent chromatin and motif evidence in Fig. 3d,e,f, this provides three complementary evaluation layers: retention of perturbation-dependent expression structure, agreement with external regulatory evidence, and model-internal sensitivity to motif perturbation.

### GenPerturb links differentiation-associated expression axes to motif-level regulatory signals

We next applied GenPerturb to lineage-associated perturbation responses, focusing on erythroid differentiation in the Norman K562 CRISPRa dataset (Fig. 4) [20]. We used perturbation program annotations and erythroid signature gene sets from the original study to evaluate whether lineage-associated structure was retained in the fitted expression profiles. Perturbations annotated as erythroid, granulocyte, megakaryocytic, and cell-cycle related programs formed structured groups in both observed and fitted low-dimensional expression spaces, indicating that multiple annotated biological axes were preserved in the model-derived profiles (Fig. 4a). The erythroid axis was then quantified using the corresponding signature gene set. Signature scores computed from fitted profiles were highly concordant with those derived from observed data, and erythroid-promoting perturbations retained high fitted scores. These results indicate that the fitted sequence-based outputs preserve a biologically interpretable erythroid differentiation axis (Fig. 4b).

We next compared TF-MoDISco-derived motif significance scores from differential attribution profiles, defined as -log10-transformed q-values (-log10 q), with TF mRNA associations along the erythroid axis. Notably, many perturbations annotated as erythroid-promoting in the original study correspond to non-transcription factor genes rather than direct regulators (Fig. 4a,b). Despite this indirect perturbation setting, GenPerturb successfully recovered canonical erythroid transcription factors such as GATA1, KLF1, and TAL1 as top-associated regulators along the erythroid axis. This suggests that the model captures downstream regulatory structure rather than merely reflecting the perturbed gene identities.

Regulators such as KLF1, TAL1, and NFE2 showed concordant positive associations in both layers, consistent with established erythroid transcription factor networks [28]. However, associations based on motif significance scores did not simply mirror TF mRNA associations across regulators, and GATA1 signals were prioritized more strongly by motif significance scores than would be expected from TF mRNA association alone. This pattern supports the interpretation that the motif significance layer captures a regulatory signal that is not fully reflected by TF mRNA abundance alone (Fig. 4c). In addition, motif degeneracy within TF families means that attribution at a single motif instance may reflect contributions from multiple related regulators rather than a one-to-one correspondence with individual TF expression trajectories. We then performed in silico motif mutation to test whether lineage-associated motifs contributed to the fitted erythroid signature shifts within the model. Randomizing prioritized motif instances attenuated erythroid signature contributions in lineage-consistent directions, supporting a model-internal link between the motif significance signal and differentiation-associated fitted outputs (Fig. 4d).

Similar analyses along the granulocyte axis recovered CEBP- and SPI1/PU.1-related motif significance signals as the top-ranked programs, consistent with their established role as lineage-determining regulators in granulocyte-macrophage regulatory programs [29]. In silico mutation of these motif instances attenuated the fitted granulocyte signature in lineage-consistent directions, in contrast to the erythroid axis where GATA motifs dominated. Together, these results indicate that the same framework can resolve distinct lineage-associated regulatory programs within the same dataset and prioritize axis-specific TF families through motif significance scores rather than recovering a generic differentiation signal (Fig. S11).

To contextualize these lineage-associated signals relative to existing regulatory inference approaches, GenPerturb-derived regulator rankings along the erythroid and granulocyte axes were compared with motif enrichment over GenPerturb regions, rE2G extended, rE2G, ABC, TSS-proximal regions, and SCENIC regulon activity [6]. These methods recovered overlapping but non-identical regulator sets, including GATA, KLF, TAL, NFE, CEBP, and SPI1-related factors, supporting a complementary interpretation of sequence attribution, regulon activity, and enhancer annotation rather than a strict ranking among methods (Fig. S12).

### Attribution-derived motif significance scores capture compound-associated regulatory activity beyond TF expression

We next analyzed compound perturbations in the Jiang et al. CD8 T-cell dataset to test whether GenPerturb-derived motif significance scores could recover upstream regulatory programs activated by compound treatment, including cases where TF activity is controlled through ligand binding, protein stability, or nuclear translocation rather than TF transcription [22].

As a positive-control setting, we first examined glucocorticoids, which bind and activate the glucocorticoid receptor NR3C1 and were represented by multiple corticosteroid compounds in the dataset [30]. Upon ligand binding, NR3C1 is released from cytoplasmic chaperone complexes, translocates to the nucleus, and activates glucocorticoid response elements without requiring increased NR3C1 transcription [30–31]. NR3C1-associated motifs were significantly enriched among glucocorticoid-treated samples, consistent with activation of glucocorticoid receptor regulatory programs at the sequence-attribution level (Fig. 5a). Notably, NR3C1 expression was not consistently induced across these compounds, indicating that motif significance scores capture regulatory activity beyond TF transcript abundance alone (Fig. 5b).

**Figure 5.**
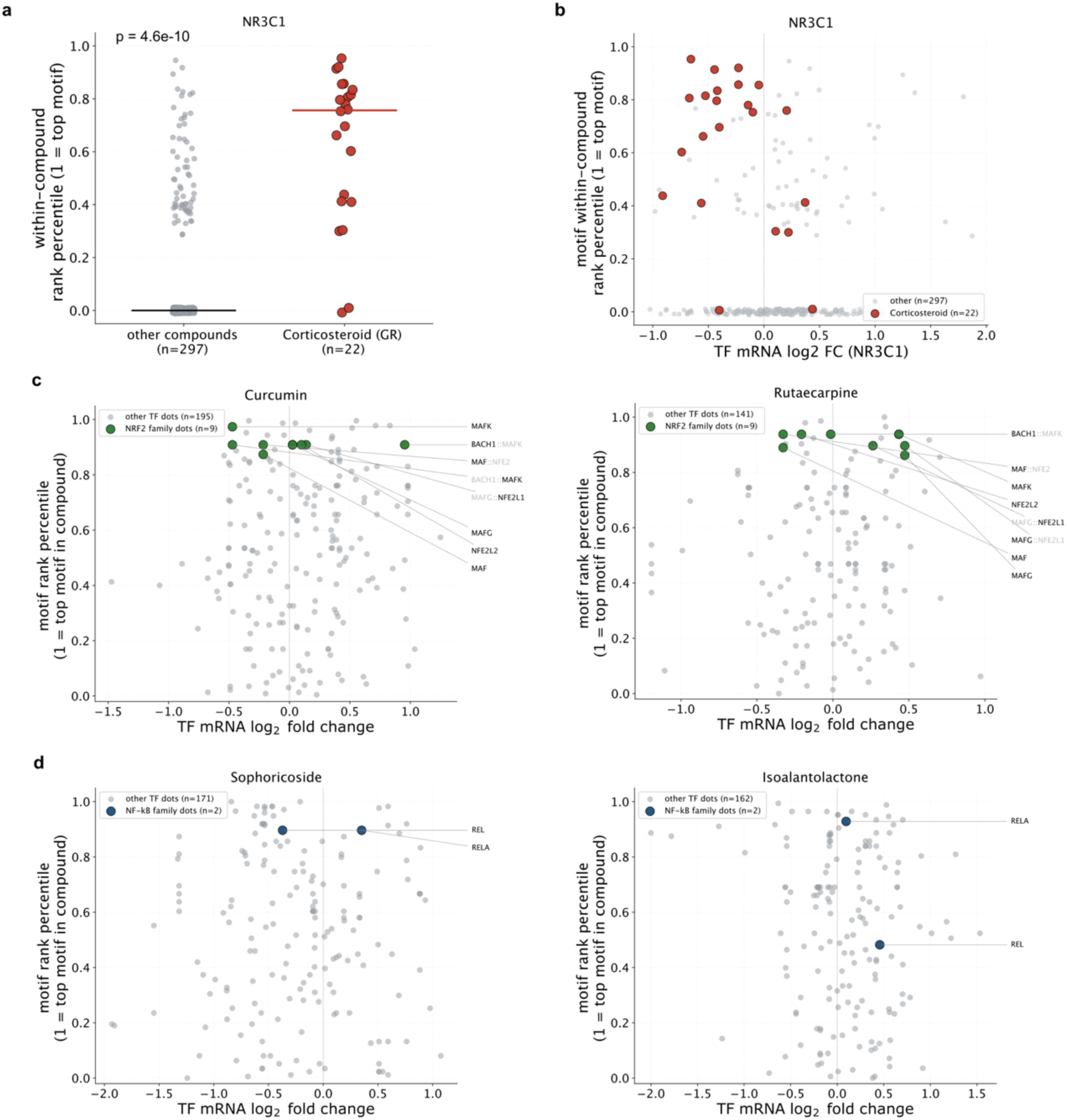
Attribution-derived motif significance scores recover activity-like regulatory programs in compound perturbations. GenPerturb was applied to the Jiang et al. CD8 T-cell compound Perturb-seq dataset to compare TF-MoDISco-derived motif significance scores (-log10 q) with TF mRNA abundance across compound perturbations. Motif significance scores are summarized as within-compound rank percentiles, with higher values indicating stronger motif significance within that compound. a, Positive-control comparison of NR3C1 motif rank percentile between corticosteroid compounds (n = 22 single-compound perturbations) and all other compounds. Each dot is a compound; horizontal bars denote group medians (one-sided Wilcoxon rank-sum p-value). b, Across-compound scatter of NR3C1 motif rank percentile (y-axis) against NR3C1 TF mRNA log2 fold change versus control (x-axis), with corticosteroid compounds highlighted in red. c, Per-compound comparison of TF mRNA change and motif rank percentile for two natural products selected to probe NRF2 (NFE2L2)-linked stress-response programs, Curcumin and Rutaecarpine. Each dot represents a motif-associated TF, with heterodimer motifs represented by their component TFs. For heterodimer motif labels, the component TF whose mRNA value is plotted is shown in black, matching monomer labels, whereas the partner TF not used for the mRNA coordinate is shown in light gray. NRF2-family motifs are highlighted in green; other TF dots are shown in gray. d, Same per-compound layout as panel c for two natural products with reported NF-κB-inhibitory anti-inflammatory activity, Sophoricoside and Isoalantolactone, using the same heterodimer label-color convention. NF-κB-family motifs (REL, RELA) are highlighted in dark blue; other TF dots are shown in gray. Both panels c and d show that target-class motifs appear within the top motif rank percentile range even when the corresponding TF mRNA is not strongly induced or repressed, consistent with model-internal attribution capturing upstream regulatory state that is not directly reflected in TF transcript abundance.

We next examined natural-product compounds, where phenotypic screens often identify bioactivity before the upstream regulatory mechanism is fully resolved. Curcumin and Rutaecarpine both showed strong enrichment of NRF2-associated motifs despite limited induction of NRF2 expression. This is consistent with known regulation of NRF2/NFE2L2 activity through KEAP1-dependent degradation and stress-induced protein stabilization, rather than transcriptional induction of the TF [32]. Curcumin has been reported to stabilize NRF2 protein through KEAP1 cysteine modification without increasing Nrf2 mRNA, and Rutaecarpine has been reported to inhibit KEAP1-NRF2 interaction and promote NRF2 activation (Fig. 5c) [33–34]. We similarly examined Sophoricoside and Isoalantolactone, compounds with reported anti-inflammatory effects mediated in part through inhibition of NF-κB signaling, to test whether their attribution profiles implicated NF-κB-associated regulatory programs. Both compounds showed RELA-associated motif signals without strong induction of REL/RELA mRNA. Because RELA/p65 activity is primarily regulated through IκBα/NFKBIA-dependent cytoplasmic retention, signal-dependent IκBα degradation, and nuclear translocation rather than by RELA transcript abundance, this result suggests that motif significance scores can identify NF-κB-linked regulatory responses even when TF mRNA levels are not strongly induced (Fig. 5d) [35–39].

Together, these examples show that GenPerturb-derived motif significance scores can recover regulatory programs linked to compound classes whose activity is mediated by ligand-dependent receptor activation, KEAP1–NRF2 protein stabilization, or NF-κB-linked regulatory state changes. These results support sequence-level motif significance scoring as a complementary framework for compound mechanism-of-action analysis, particularly in phenotypic screens where TF mRNA abundance alone may fail to identify the relevant upstream regulatory program. We present these compounds as illustrative positive-control and candidate examples rather than as a compound-wide systematic benchmark.

## Discussion

In this study, we introduced GenPerturb, a model-based interpretation framework that converts gene-level perturbation responses into element-level, locus-anchored cis-regulatory hypotheses using pretrained sequence representations. By fitting perturbation-resolved expression outputs from a shared genomic input and comparing attribution profiles between control and perturbation states, GenPerturb prioritizes candidate regulatory regions, motif instances, and perturbation-associated regulatory programs from Perturb-seq data (Fig. 1). The main contribution is the translation of gene-level perturbation responses into locus-resolved, sequence-grounded regulatory hypotheses.

This framework addresses a central limitation of standard Perturb-seq analysis [1–2]. While Perturb-seq captures downstream transcriptional responses, it does not directly resolve the underlying cis-regulatory elements or sequence motifs. Conventional approaches typically rely on post hoc analyses, such as differential expression, pathway enrichment, and motif enrichment [3–8]. In contrast, GenPerturb directly links perturbation responses to regulatory sequence features within a shared sequence-based framework, enabling interpretation at the level of candidate regulatory regions and motif instances. Rather than asking which motifs are enriched in responsive gene sets, GenPerturb identifies motif instances at specific loci associated with perturbation responses.

Our results support three main conclusions. First, pretrained sequence representations can be transferred to Perturb-seq datasets and retain meaningful perturbation-dependent expression structure across CRISPR activation, interference, knockout, and compound perturbation datasets. Comparisons with Enformer, Borzoi, AlphaGenome, and a Simple CNN baseline highlight the importance of pretrained regulatory features [13–15]. Notably, alignment to the target assay and cell state requires dataset-specific fitting, indicating that pretrained outputs alone are insufficient. Thus, GenPerturb uses foundation sequence models as transferable regulatory representations for interpreting perturbation responses, rather than as standalone predictors.

Second, control-subtracted differential attribution enables prioritization of perturbation-associated cis-regulatory regions and motif instances supported by independent chromatin information. A key aspect is the comparison between fitted control and perturbation outputs on the same genomic input, which mitigates constitutive contributions and highlights perturbation-specific effects. Using this approach, GenPerturb showed a small but consistent advantage over rE2G extended, rE2G, ABC, and TSS-distance baselines, particularly for distal regulatory elements (Fig. 3d) [4–5]. Rather than indicating broadly superior enhancer prediction, these results support the idea that an RNA-only sequence-fitted model can recover biologically meaningful signals. Unlike annotation-based approaches that rely on precomputed enhancer–gene links, GenPerturb is fitted directly to the perturbation dataset and identifies sequence features whose contribution changes between conditions. Motif analyses similarly demonstrate improved recovery compared with annotation-derived regions [3,7–8], and in silico motif perturbation attenuates fitted responses, supporting a link between prioritized sequence features and perturbation-associated outputs (Fig. 3c,f). Together, these properties position GenPerturb as complementary to existing enhancer-linking frameworks and suitable for prioritizing regulatory hypotheses.

Third, applications to differentiation and compound perturbation show that motif-level attribution captures regulatory signals not fully explained by transcription factor mRNA abundance. In lineage differentiation, GenPerturb recovered known regulatory programs, including cases where TF transcripts showed minimal changes. In compound perturbations, motif programs consistent with NR3C1, NRF2, and NF-κB signaling were identified even when corresponding transcripts changed little (Fig. 5) [30,32–33,35–37]. Many compounds modulate TF activity without corresponding transcript-level shifts, limiting inference based on TF mRNA alone. By relying on sequence-based attribution, GenPerturb provides complementary regulatory insight and supports hypothesis generation for mechanism-of-action studies.

More broadly, GenPerturb illustrates how foundation sequence models can be integrated into perturbation biology as an interpretation framework. By combining sequence-to-expression modeling with differential attribution, it links perturbation responses to specific regulatory regions and motif instances in a unified framework, bridging sequence modeling, enhancer annotation, and regulon inference. This approach complements multiomic regulatory inference methods and differs from observational sequence-to-expression models by explicitly contrasting fitted control and perturbation outputs [40–43]. It is particularly useful for retrospective analysis of RNA-only Perturb-seq datasets lacking matched chromatin data.

Several limitations should be considered. First, GenPerturb is fitted to observed perturbation conditions within a specific cell type and is not designed to predict responses to unseen perturbations or cell types, but rather to interpret responses within a defined experimental context [44]. Second, attribution reflects model-derived interpretation rather than direct measurement of enhancer activity, transcription factor binding, chromatin accessibility, or protein-level regulatory activity; prioritized regions and motif instances should therefore be regarded as hypotheses requiring experimental validation [45]. This is particularly relevant in compound perturbation settings, where motif-level signals can nominate candidate regulatory programs but do not by themselves establish targets or mechanisms of action. Third, motif interpretation is constrained by motif degeneracy and transcription factor family ambiguity, limiting unique TF assignment. Fourth, attribution depends on the pretrained sequence representation, which may introduce biases; in practice, the backbone is typically frozen, making sequence sensitivity largely determined by the pretrained model, with perturbation specificity captured in condition-dependent outputs. Fifth, the current framework operates on condition-level pseudobulk outputs and does not resolve cellular heterogeneity. Sixth, limited availability of perturbation-resolved multiome datasets constrains external validation, with benchmarking largely anchored to available resources such as the Martin et al. erythroid dataset (Supplementary Table 3). Finally, the reliability of attribution depends on the quality of the fitted perturbation response, and weak or noisy signals may reduce interpretability.

In summary, GenPerturb provides a practical framework for extracting sequence-grounded regulatory hypotheses from RNA-only perturbation data by integrating pretrained sequence representations, perturbation-resolved modeling, and differential attribution.

## Conclusions

GenPerturb translates gene-level perturbation responses into locus-resolved, sequence-grounded regulatory hypotheses by fitting pretrained sequence-to-expression representations to observed Perturb-seq outputs and contrasting attribution profiles between control and perturbation states. This approach captures perturbation-dependent expression structure and prioritizes regulatory regions and motif instances associated with perturbation responses, including regulatory programs not fully explained by transcription factor expression changes. By operating directly on RNA-only perturbation datasets without requiring matched chromatin measurements, GenPerturb complements existing regulatory inference approaches and provides a unified, sequence-anchored view of genes, regulatory elements, and motif instances. It serves as a practical hypothesis-prioritization layer for both prospective and retrospective Perturb-seq studies, enabling downstream experimental validation of candidate regulatory mechanisms.

## Methods

### Data preprocessing for perturbation data

We assembled twelve perturbation datasets from five published studies covering CRISPR activation, CRISPR interference, CRISPR knockout, and small-molecule perturbations: Norman et al. K562 CRISPRa [20], Replogle et al. K562 genome-wide and essential-gene CRISPRi screens and an RPE1 essential-gene CRISPRi screen [21], Martin et al. erythroid-lineage Perturb-multiome [12], Jiang et al. compound perturbations in primary CD4 T, CD8 T, B, and myeloid cells [22], and Srivatsan et al. sci-Plex compound perturbations in A549, K562, and MCF7 [23]. Processed single-cell objects for Norman, Replogle, and Srivatsan were obtained from scPerturb [46] and the sc-pert repository (https://github.com/theislab/sc-pert). The Jiang et al. datasets were downloaded from the original study repository (GSE217460, https://data.caltech.edu/records/2cjss-wgh69). The Martin et al. erythroid Perturb-multiome data were downloaded from GEO (GSE274113).

For CRISPR datasets, unperturbed cells (those that failed gRNA-induced expression change of the target gene) were excluded using Mixscape [47] as implemented in pertpy (v0.10.0) [48]. Highly variable genes were defined within each gRNA cell group using ‘scanpy.pp.highly_variable_genes‘ with ‘min_disp = 0.2‘ (Scanpy v1.11.2) [49], and the union of HVGs across gRNA groups was retained.

For all datasets, Pseudobulk expression matrices were generated using adpbulk (v0.1.4) with ‘method = "sum"‘, summing UMI counts per perturbation condition. Perturbation conditions represented by 100 cells or fewer after Mixscape filtering were removed to ensure stable condition-level expression estimates. Pseudobulk counts were CPM-normalized and log2(x + 1)-transformed, and only genes with CPM > 2 in at least one perturbation condition were retained.

For knockdown (CRISPRi) and overexpression (CRISPRa) perturbations targeting expressed genes, the pseudobulk expression value of the target gene itself was replaced with the corresponding value from the non-targeting control before model fitting, preventing the model from learning a trivial sequence-to-expression mapping in which the target gene’s own promoter directly encodes its perturbation response. Martin et al. erythroid Perturb-multiome dataset was processed through the same Mixscape, HVG, pseudobulking, CPM normalization, log2 transform, and gene-filter steps. For the matched ATAC-seq modality, cells passing the RNA-seq–based filtering and perturbation-label assignment steps were retained, and ATAC profiles were pseudobulked using the corresponding perturbation-condition labels.

Control conditions were defined according to each original study: non-targeting gRNA for Norman, Replogle, and Martin et al.; the drug-free control condition for Jiang et al.; and vehicle (DMSO) for Srivatsan et al.

### Reference genome and training/validation/test split

The human reference genome was GRCh38.p14, and gene models were obtained from the GENCODE v46 basic annotation [50]. Only protein-coding mRNAs and lncRNAs were retained. For each gene, one representative transcription start site (TSS) was selected by prioritizing transcripts annotated as Ensembl canonical; when absent, transcripts were ordered by annotation rank (as provided in the GENCODE annotation), with transcript identifiers serving as a deterministic tie-breaker.

Training, validation, and test partitions followed the genome partitioning scheme defined in the AlphaGenome pretraining dataset [15]. Fold assignments were obtained from the official AlphaGenome ‘fold_intervals‘ interface and mapped to the retained representative TSS positions. All evaluations reported here were conducted using a fixed downstream split corresponding to fold_0. This same downstream partition was consistently applied across all backbone models (AlphaGenome, Enformer, Borzoi, and a Simple CNN baseline) to ensure fair comparison.

### DNA foundation model embeddings

To decouple representation extraction from prediction-head training, embeddings from pretrained DNA foundation models were computed once for every retained representative TSS and cached for downstream use.

1. AlphaGenome Embeddings were extracted with the ‘alphagenome-pytorch‘ re-implementation (v0.2.6) [15]. The official ‘alphagenomè Python package was used to obtain genome-fold intervals. The primary transfer models used embeddings from the all-fold AlphaGenome checkpoint. A context window of 1,048,576 bp centered on each TSS was provided to the model, and activations were taken from the layer immediately preceding the final prediction head. The four 128-bp bins (512 bp in total) surrounding the TSS were retained directly, yielding a per-gene representation of shape (4, 3072) = 12,288 dimensions, where 3072 is the number of channels in the extracted hidden representation.
2. Enformer. Enformer embeddings were extracted with the ‘enformer-pytorch‘ implementation (v0.8.10) [13]. A context window of 196,608 bp centered on each TSS was used, and the four 128-bp bins surrounding the TSS were retained directly to produce a per-gene representation of shape (4, 3072) = 12,288 dimensions, where 3072 is the number of channels in the extracted hidden representation.
3. Borzoi. Borzoi embeddings were extracted with the ‘borzoi-pytorch‘ implementation (v0.4.2) [14]. A context window of 524,288 bp centered on each TSS was provided to the model. At Borzoi’s 32-bp resolution, 16 bins (≈512 bp) surrounding the TSS were retained directly, yielding a per-gene representation of shape (16, 1920) = 30,720 dimensions, where 1920 is the number of channels in the extracted hidden representation.
4. Simple CNN baseline. A non-pretrained baseline operating on 40,001-bp DNA windows centered on the TSS was trained end-to-end from random initialization without prior representation learning.

### Model architecture

For all backbones the prediction head was a multi-output linear layer mapping the per-gene embedding to a vector with one element per perturbation condition, followed by a ReLU activation. The output dimensionality equals the number of perturbation conditions in the dataset, comprising every perturbation together with the non-targeting control, and output values correspond to per-gene log2(CPM + 1) expression for each condition. Importantly, perturbation identity is not provided as an input feature; perturbation-dependent responses are represented entirely as differences among the condition-specific output channels anchored to the same gene locus and sequence coordinate. The same head architecture was used across AlphaGenome, Enformer, Borzoi, and the Simple CNN baseline, differing only in input dimensionality.

### Training

All training was performed with PyTorch v2.3.1 (PyTorch v2.4.1 + CUDA 12.1 in the Borzoi environment) using NVIDIA H200 GPUs [51]. The optimiser was AdamW with learning rate 1 × 10⁻⁴ and weight decay 5 × 10⁻³, the loss was mean squared error (MSE), and the gradient clipping value was 0.2. Training was terminated when the validation MSE did not improve for twenty consecutive epochs. Four training strategies were used:

1. **Feature extraction (transfer).** Pre-computed embeddings were fed directly to the prediction head; the backbone was held frozen. Training ran for up to 100 epochs with batch size 256 in fp32 precision. This was the default strategy and was used throughout the main results.
2. **LoRA fine-tuning.** Low-rank adaptation [25] was applied through the PEFT library (v0.4.0) [52]. Adaptation targeted the attention query, key, value, and output projections of the backbone (‘to_qkv‘ and ‘to_out‘ for AlphaGenome) while the prediction head remained trainable, with α = 2 and rank r ∈ {64, 256, 512}. Training ran for up to 100 epochs at effective batch size 256 (per-device batch 2 with 128-step gradient accumulation) in bf16 mixed precision.
3. **Full fine-tuning.** The entire backbone together with the head was fine-tuned for up to 150 epochs at the same effective batch size of 256 in bf16 mixed precision.
4. **Simple CNN baseline.** The Simple CNN was trained end-to-end from random initialization on 40,001-bp DNA sequence input for up to 150 epochs at effective batch size 256 in bf16 mixed precision.

For AlphaGenome cached-embedding extraction and sequence-mode runs (LoRA, full fine-tuning, and DNA-side prediction), the running-variance update of the internal ‘BatchRMSNorm‘ modules was disabled to prevent evaluation-time statistics drift. For AlphaGenome sequence-mode runs, gradient checkpointing was enabled in the transformer and U-Net blocks to fit the 1,048,576-bp context window into memory. For feature-extraction runs, predictions were obtained by applying the trained head to the cached per-gene embedding. For fine-tuning, LoRA, and Simple CNN runs, predictions were obtained by forwarding the representative-TSS-centered sequence through the backbone using the GRCh38.p14 reference.

### Pearson correlation evaluation

Agreement between observed and fitted outputs was assessed on the held-out test split using two complementary Pearson correlations. The *across-genes* correlation was computed per perturbation between observed and fitted values, summarizing how well the model reproduces the gene-level expression profile of each condition. The *across-perturbations* correlation was computed per gene between observed and fitted log2(CPM + 1) values across all perturbation conditions, capturing condition-dependent variation at a fixed locus. Genes were stratified into five quintile bins of mean expression to expose expression-level effects on correlation (Fig. 2a,c).

### Pretrained versus transferred AlphaGenome under unperturbed conditions

To test whether the original AlphaGenome representation could be used directly as an expression output, raw AlphaGenome predictions under the unperturbed condition were contrasted with predictions from the transferred model in CD8T and K562 cells. AlphaGenome raw outputs were taken from the prediction-head layer that concatenates the seven human genome track heads (rna_seq, cage, dnase, procap, atac, chip_tf, chip_histone) into 4,992 tracks; for each dataset, CAGE-seq replicates corresponding to the study cell types (CD8T or K562) were identified using the published track metadata, summed across the four 128-bp center bins surrounding the TSS and across the ± strand pair, and log-transformed before comparison with the observed log2(CPM + 1) values and with the transferred AlphaGenome prediction.

### Baseline predictors

To assess the contribution of perturbation-specific fitting beyond trivial expression structure, two baseline predictors were constructed. The control-copy baseline replaced values for all non-control perturbations with those of the control condition plus small Gaussian noise (σ = 0.001). The perturbation-mean baseline replaced each perturbation with the gene-wise mean across all perturbations (excluding the control) plus identical noise, while the control condition was preserved. Baselines were evaluated using the same across-gene and across-perturbation Pearson correlation pipeline as the trained models.

### Embedding and clustering analysis

Observed and fitted expression matrices were each loaded as Scanpy v1.11.2 AnnData objects (rows = perturbation conditions; columns = genes), with genes z-scored using ‘sc.pp.scalè. PCA, a k-nearest-neighbor graph (‘n_neighbors = 10‘), Leiden community detection (‘resolution = 1.5‘), and UMAP were computed independently on observed and fitted profiles restricted to the held-out test set. To visualize cluster transfer between modalities, observed Leiden labels were overlaid on the fitted UMAP, producing the paired observed/fitted UMAP panels in Fig. 2d, Fig. S3a, and Fig. S5a,b.

### Cluster concordance metrics

Leiden clustering was performed independently on the observed and fitted expression matrices, and concordance between the resulting partitions was quantified using the adjusted Rand index (ARI) and normalized mutual information (NMI) computed with scikit-learn (v1.7.0).

### Multifold evaluation

Robustness to the train/validation/test partition was assessed using the four rotating split configurations defined by the AlphaGenome pretraining scheme (fold_0–fold_3). In each fold, both the genomic data partition and the backbone embeddings were fold-specific: the transferred prediction head was trained, validated, and tested on that fold’s own train/validation/test split, using embeddings from the matching pretrained AlphaGenome checkpoint.

We additionally included an "all_folds" condition, in which embeddings were drawn from the AlphaGenome all_folds checkpoint. In this setting, the downstream prediction head was trained, validated, and tested on the fold_0 data split (i.e., the same downstream partition as the fold_0 setting).

### Differential attribution and peak calling

For track visualization, motif discovery, and in silico mutation analyses, per-base attribution scores were computed using Captum (v0.8.0) [53]. For each selected input sequence, Input × Gradient [16] and Saliency [54] were computed for the non-targeting control output channel and for the target perturbation output channel. The perturbation-specific differential attribution track was defined as the element-wise difference between perturbation-channel and control-channel Input × Gradient on the same input sequence. For peak calling, the per-base differential attribution scores were first converted to absolute values and then binned to 128-bp resolution. The resulting 128-bp binned absolute attribution tracks were processed with MACS3 bdgpeakcall (v3.0.3) [55] using cutoff = 0.0001, minimum length = 100 bp, and maximum gap = 150 bp. The 128-bp windows overlapping each called peak were retained for motif discovery and in silico mutation analyses. For visualization, signed differential attribution tracks were also written as BedGraph files.

### Gene-level evaluation of raw and differential attribution against observed fold changes

For the Martin et al. dataset, raw output-channel attribution and control-subtracted differential attribution were compared at the gene level using observed fold changes as the reference measure (Fig. S8). For each perturbation with genome-wide attribution profiles, three per-gene attribution summaries were computed:

1. the sum of absolute Input × Gradient attribution for the control output
2. the sum of absolute Input × Gradient attribution for the perturbation output
3. the sum of absolute control-subtracted differential Input × Gradient attribution Attribution scores were matched to pseudobulk |FC| values for the same genes, where |FC| was computed as the absolute difference in log2(CPM + 1) between each perturbation and the non-targeting control.

Rank-based enrichment was evaluated for perturbations with at least 100 evaluable genes. Within each perturbation and attribution type, genes were ranked independently by attribution magnitude and observed |FC|. Top-ranked gene sets were selected using matched percentile cutoffs ranging from 0.25% to 25% of evaluable genes. For each cutoff, the overlap between attribution-ranked and |FC|-ranked genes was divided by the overlap expected under independence, k²/n, where k is the number of selected genes and n is the number of evaluable genes. Enrichment ratios were then averaged across perturbations.

Gene-wise consistency across perturbations was evaluated by computing, for each gene, the Spearman correlation between attribution magnitude and observed |FC| across perturbations. This analysis was performed separately for raw perturbation-output attribution and control-subtracted differential attribution. Genes were further grouped by their maximum observed |FC| across perturbations to assess how attribution–response agreement varied with response magnitude.

### Enhancer AUPRC benchmark

In the Martin et al. benchmark, the RNA modality was used as the target response for model fitting. Matched ATAC-derived TF perturbation–responsive accessible chromatin regions from Table S2 of Martin et al. and TF perturbation–responsive differentially expressed genes (DEGs) from Table S3 of the same study were used only for benchmark construction.

TF perturbation–responsive accessible chromatin regions from Martin et al. Table S2 were converted into per-perturbation reference BED files. Candidate regions were constructed for genes listed in the Martin et al. TF perturbation–responsive DEG table (Table S3) with |log2FC| ≥ 0.50 by combining GenPerturb attribution peaks with precomputed rE2G extended, rE2G, and ABC region sets provided as BED files [4–5]. Because cell-context-matched rE2G and ABC predictions were not available for the Martin et al. erythroid context, K562 predictions were used as the closest available erythroid-lineage proxy.

All source-specific BED files were restricted to the 1,048,576-bp TSS-centered model input window for each target gene. Regions from GenPerturb, rE2G extended, rE2G, ABC, and the TSS-distance baseline were merged within each target gene to define a shared candidate-region universe for each perturbation. Overlapping genomic intervals linked to different genes were retained as distinct enhancer–gene relationships.

Candidate regions were labelled positive if they overlapped the corresponding TF perturbation–responsive accessible chromatin region from Martin et al. Table S2 for the same perturbation; all other candidate regions were labelled negative. For each candidate region, scores were assigned from differential attribution, rE2G extended, rE2G, ABC, and TSS distance. Differential attribution scores were calculated from the raw per-base differential Input × Gradient array. At each base pair, attribution magnitude was defined as the sum of absolute values across the four nucleotide channels. The region-level attribution score was defined as the mean magnitude over the top 10% highest-scoring base pairs within the candidate region. ENCODE-rE2G, ENCODE-rE2G extended, and ABC scores were joined only from overlapping records linked to the same target gene. When multiple matching records overlapped a candidate region, the maximum score was used. Candidate regions without an overlapping same-gene rE2G, rE2G extended, or ABC record were retained and assigned a score of zero for that method. The TSS-distance baseline was defined as 1 / (1 + d), where d is the base-pair distance from the candidate-region midpoint to the target gene TSS. AUPRC was computed separately for each perturbation and for four TSS-distance strata (0–1 kb, 1–10 kb, 10–100 kb, and >100 kb). Perturbation–stratum combinations with fewer than 10 positive regions were excluded. Mean AUPRC across perturbations, paired comparisons between methods within each distance stratum, and within-perturbation 95% bootstrap confidence intervals were calculated using 1,000 bootstrap iterations.

### Motif discovery and recovery

TF-MoDISco analysis was performed with tfmodisco-lite (v2.5.2) [26] on differential attribution profiles. For each perturbation, the 200 genes with the largest absolute observed fold changes and their 128-bp attribution peak windows were used as input. Saliency was supplied as the hypothetical-contribution input after per-position zero-centering across the four nucleotide channels, following the hypothetical-contribution convention used by tfmodisco-lite, and tfmodisco-lite internally constructed the contribution signal as the element-wise product of the one-hot reference sequence and this hypothetical contribution. Parameters were ‘sliding_window_size = 15‘, ‘flank_size = 5‘, ‘target_seqlet_fdr = 0.2‘, ‘trim_to_window_size = 15‘, ‘initial_flank_to_add = 5‘, ‘final_flank_to_add = 5‘, ‘final_min_cluster_size = 20‘, and ‘subcluster_perplexity = 10‘. Discovered patterns were annotated with Tomtom (MEME Suite v5.5.9) [7–8] against the JASPAR 2022 CORE vertebrate non-redundant PFM database [27]. Complementary known-motif enrichment was performed with GimmeMotifs (v0.18.1) [3] using ‘gimme motifs --known‘ against the JASPAR 2022 vertebrate PFM database, applying the same enrichment procedure to attribution, rE2G extended, rE2G standard, ABC, and TSS ± 1 kbp region sets. Per-perturbation TF-MoDISco Tomtom matches and GimmeMotifs motif matches were summarized at q < 0.05 as gene matches to the perturbed TF and as JASPAR-cluster matches to the perturbed TF family.

### In silico motif mutation and representative locus

Motif regions were defined using the width of JASPAR motifs matched to TF-MoDISco–identified motifs. For the Fig. 3f and Fig. S10b boxplots, we tested whether model-internal sequence sensitivity increased with differential-attribution strength using motifs whose Tomtom annotation matched the perturbed TF or a TF in the same JASPAR cluster at q < 0.05. Every unique matched site was retained (2,735 Martin et al. sites across 515 genes; 8,323 Norman sites across 1,226 genes). For each motif, a length-matched negative-control interval was placed at the nearest available offset from a discrete set of {±500, ±1,000, ±2,000, ±3,000} bp within the 1,048,576-bp model input window and outside annotated motif regions extended by 10 bp. Each motif target and paired negative-control interval was mutated five times with independent mononucleotide-preserving random shuffles of the reference bases, and all perturbation output channels were re-predicted for every shuffled sequence with the trained GenPerturb model. The choice of mononucleotide-preserving shuffling as the in-silico motif-perturbation control follows the convention adopted by scooby [42]. The five mutant predictions were averaged for each target before calculating downstream mutation effects. The cancellation magnitude for each target and perturbation channel was then defined as |(seed-averaged mutant perturbation-control fold change) - (wild-type perturbation-control fold change)|.

Matched motif targets were stratified into tertiles by the absolute Input × Gradient attribution sum over the motif and compared with the paired negative-control group; High-attribution versus negative-control separation was tested with a one-sided Mann-Whitney U statistic.

### Analysis of lineage-associated perturbation programs in the Norman et al. dataset

Observed and fitted perturbation-level expression profiles from the Norman et al. K562 CRISPRa dataset were projected into two-dimensional space using UMAP and annotated with the perturbation program labels reported in the original study.

Erythroid and granulocytic perturbation programs and the corresponding signature gene sets were obtained from Norman et al. [20]. The erythroid signature comprised HBG1, HBG2, HBZ, HBA1, HBA2, GYPA, and ERMAP, whereas the granulocyte signature comprised ITGAM, CSF3R, LST1, and CD33. Signature scores were computed independently for observed and fitted AnnData objects using ‘scanpy.tl.score_genes‘ with default parameters. For each lineage signature, observed and fitted per-perturbation scores were compared across all perturbations using Pearson correlation.

### Motif versus TF mRNA correlations along the signature axis

To compare motif-level and TF expression-level associations with lineage-signature variation, we computed motif–signature and TF mRNA–signature correlations for matched motif–TF pairs. TF-MoDISco Tomtom annotations were converted into perturbation-level motif score matrices by transforming Tomtom q-values as −log10(q). For each lineage signature, Pearson correlations were calculated between motif scores and observed perturbation-level signature scores across perturbations. When multiple TF-MoDISco pattern groups were annotated to the same motif, the correlation with the largest absolute Pearson r was retained. For each matched motif–TF pair, TF mRNA expression was correlated with the same signature scores across the same perturbations. Each plotted point therefore represents the motif–signature correlation (x-axis) and the corresponding TF mRNA–signature correlation (y-axis) for a matched motif–TF pair.

### In silico motif mutation along the signature axis in the Norman et al. dataset

To test whether lineage-associated TF-MoDISco motif instances contributed to fitted signature shifts, we performed in silico motif mutation at lineage marker genes. We analyzed six erythroid marker genes (HBG1, HBG2, HBZ, HBA1, HBA2, and GYPA) and three granulocyte marker genes (ITGAM, CSF3R, and LST1). Signature genes lacking recovered TF-MoDISco motif instances (ERMAP and CD33) were excluded from the mutation analysis. Among TF-MoDISco–identified motifs at these genes, we selected motif instances belonging to the GATA, KLF, CEBP/ATF, and SP/GC-box motif families. STAT-family motifs were included as a negative-control motif class.

Only motif instances recovered by TF-MoDISco at the analyzed marker genes were considered. For each selected motif instance, the motif sequence was mutated using five independent mononucleotide-preserving random shuffles of the reference bases, following the same shuffle convention as in the Martin et al. analysis. The trained GenPerturb model was then re-evaluated on each shuffled sequence, and wild-type and mutant fold changes were calculated relative to the non-targeting control for all perturbation output channels. For each motif instance, perturbation, and marker gene, the mutation effect was defined as ΔΔpred = (mutant perturbation–control fold change) − (wild-type perturbation–control fold change) and averaged across the five shuffles. To summarize effects at the lineage-signature level, gene-level ΔΔpred values were averaged across genes within each signature.

### SCENIC regulon inference

SCENIC regulon activity per cell was inferred from the Mixscape-processed Norman AnnData using pySCENIC (v0.12.1) [6,56]. The pipeline was: (i) co-expression network inference with GRNBoost2 [57] using a human TF list as the regulator vocabulary; (ii) cisTarget motif-based pruning against the v10 clustered databases (‘hg38_10kbp_up_10kbp_down_full_tx_v10_clust‘ rankings and the ‘motifs-v10nr_clust-nr.hgnc-m0.001-o0.0‘ motif annotation); (iii) regulon construction from pruned motifs; and (iv) AUCell scoring per cell. Per-cell AUCell scores were averaged within each perturbation to obtain a per-regulon perturbation-level activity matrix.

### Lineage-regulator UpSet plots

Lineage-regulator summary plots were generated to compare how predefined lineage transcription factors (TFs) were prioritized by six regulatory evidence sources. The sources were: (i) TF-MoDISco motif scores from GenPerturb differential attribution; (ii–v) GimmeMotifs enrichment scores computed on GenPerturb attribution peaks, rE2G extended, rE2G, ABC, and TSS ± 1 kbp regions, respectively; and (vi) SCENIC regulon activity summarized by mean AUCell score per perturbation. For each source, we computed the correlation between its scores and the lineage signature score across perturbations, and ranked features by the absolute Pearson correlation coefficient (p < 0.05). For visualization, we used an UpSet-style representation in which evidence sources are arranged along the horizontal axis and predefined lineage regulators along the vertical axis. The lineage-defining TF sets were defined as follows: erythroid = {GATA1, KLF1, TAL1, LMO2, NFE2} and granulocyte = {CEBPA, SPI1, CEBPB, CEBPE, GFI1, IRF4, IRF8, BATF3, KLF4, MAFB}. Within each evidence source, all TFs (motif-family clusters) were ranked by the absolute Pearson correlation coefficient (|r|). The lower matrix summarizes the ranking of each lineage regulator, with the number inside each circle indicating its correlation-based rank. The upper bar plots show the total number of significant TF–motif clusters. Because SCENIC regulons are generated using motif-ranking databases that already group TFs with similar binding specificities, no additional collapsing into motif families was applied to SCENIC-derived results. For GimmeMotifs-based sources, motifs were retained only when at least one associated TF showed a mean observed log2(CPM) expression ≥ 0.5.

### Glucocorticoid positive-control benchmark

Analyses were performed using the Jiang et al. CD8 T-cell compound Perturb-seq dataset [22] and the corresponding fitted GenPerturb model trained with AlphaGenome feature-extraction transfer. Motifs were ranked within each compound by −log10(q-value), with non-detected motifs assigned a score of 0. NR3C1 mRNA log2 fold change relative to the non-perturbed control was calculated from observed pseudobulk counts with a pseudocount of 1. Only single-compound perturbations were included.

Single compounds annotated as corticosteroids based on PubChem and related pharmacological resources were assigned to a corticosteroid group (n = 22), and all remaining single compounds were assigned to the background group. NR3C1 motif rank percentiles were compared between groups using a one-sided Wilcoxon rank-sum test. NR3C1 motif rank percentiles were also compared with NR3C1 mRNA log2 fold changes across all single compounds.

### Per-compound motif × TF mRNA decomposition

For each compound, motifs passing the TF-MoDISco significance threshold (q ≤ 0.05) were retained and rank percentiles were recomputed within the filtered set.

Motif names were split on "::", and each component was mapped to a gene symbol by case-insensitive matching to the pseudobulk expression matrix. For heterodimer motifs, each TF component was assigned the same motif rank percentile and paired with its corresponding mRNA log2 fold change. For visualization, TF-family annotations were used to highlight NRF2-related motifs (NFE2L2, MAF, MAFB, MAFF, MAFG, MAFK, MAF::NFE2, MAFG::NFE2L1, and Bach1::Mafk) and NF-κB-

related motifs (REL and RELA).

## Data availability

The perturbed scRNA-seq data were obtained from multiple sources: Norman et al. and Replogle et al. from scPerturb (http://projects.sanderlab.org/scperturb/); Srivatsan et al. from the sc-pert repository (https://github.com/theislab/sc-pert); Jiang et al. from GSE217460 (https://data.caltech.edu/records/2cjss-wgh69); and Martin et al. from GEO (GSE274113). Martin et al. Supplementary Tables S2 and S3 associated with the original article [12] were used for TF-sensitive ACR and TF-sensitive gene annotations. The GRCh38.p14 reference genome and GENCODE v46 basic gene annotation were downloaded from GENCODE release 46 (https://www.gencodegenes.org/human/release_46.html), including ‘GRCh38.p14.genome.fa.gz‘, ‘gencode.v46.chr_patch_hapl_scaff.basic.annotation.gff3.gz‘, and ‘gencode.v46.chr_patch_hapl_scaff.basic.annotation.gtf.gz‘; representative TSS coordinates were parsed from these annotation files by prioritizing Ensembl-canonical transcripts and using deterministic fallbacks when needed. Pretrained model weights were obtained through their model-specific download interfaces, including the official AlphaGenome JAX weights and Hugging Face repositories used by the PyTorch implementations. ENCODE-rE2G enhancer-gene predictions were downloaded from the ENCODE Portal, including the K562 standard files ENCFF497HEA and ENCFF246ZQE and K562 extended files ENCFF269DKY and ENCFF950FTI (https://www.encodeproject.org/). ABC predictions were downloaded from the Engreitz Lab AvgHiC ABC0.015 minus150 ForABCPaperV3 release (ftp://ftp.broadinstitute.org/outgoing/lincRNA/ABC/AllPredictions.AvgHiC.ABC0.015. minus150.ForABCPaperV3.txt.gz) and converted from hg19 to hg38 using the hg19-to-hg38 mapping invoked by ‘pyliftover.LiftOver("hg19", "hg38")‘, corresponding to the UCSC hg19ToHg38 chain resource (http://hgdownload.soe.ucsc.edu/goldenPath/hg19/liftOver/hg19ToHg38.over.chain.g z). Known motif sequences and cluster annotations were downloaded from JASPAR (https://jaspar.elixir.no/). SCENIC/cisTarget reference data were downloaded from the Aerts Lab resources: the human TF list (https://resources.aertslab.org/cistarget/tf_lists/allTFs_hg38.txt), the hg38 v10 clustered rankings database (https://resources.aertslab.org/cistarget/databases/homo_sapiens/hg38/refseq_r80/ mc_v10_clust/gene_based/hg38_10kbp_up_10kbp_down_full_tx_v10_clust.genes_v s_motifs.rankings.feather), and the motif-to-TF annotation table (https://resources.aertslab.org/cistarget/motif2tf/motifs-v10nr_clust-nr.hgnc-m0.001-o0.0.tbl). Compound annotations for Fig. 5 were curated from PubChem using PUG-REST and PUG-View (https://pubchem.ncbi.nlm.nih.gov/); the PubChem screening script and cached evidence tables are provided with the analysis code.

## Code availability

The analysis code is available at https://github.com/rikenbit/GenPerturb.

## Data availability

The perturbed scRNA-seq data were obtained from multiple sources: Norman et al. and Replogle et al. from scPerturb (http://projects.sanderlab.org/scperturb/); Srivatsan et al. from the sc-pert repository (https://github.com/theislab/sc-pert); Jiang et al. from GSE217460 (https://data.caltech.edu/records/2cjss-wgh69); and Martin et al. from GEO (GSE274113). Martin et al. Supplementary Tables S2 and S3 associated with the original article [19] were used for TF-sensitive ACR and TF-sensitive gene annotations. The GRCh38.p14 reference genome and GENCODE v46 basic gene annotation were downloaded from GENCODE release 46 (https://www.gencodegenes.org/human/release_46.html), including ‘GRCh38.p14.genome.fa.gz‘, ‘gencode.v46.chr_patch_hapl_scaff.basic.annotation.gff3.gz‘, and ‘gencode.v46.chr_patch_hapl_scaff.basic.annotation.gtf.gz‘; representative TSS coordinates were parsed from these annotation files by prioritizing Ensembl-canonical transcripts and using deterministic fallbacks when needed. Pretrained model weights were obtained through their model-specific download interfaces, including the official AlphaGenome JAX weights and Hugging Face repositories used by the PyTorch implementations. ENCODE-rE2G enhancer-gene predictions were downloaded from the ENCODE Portal, including the K562 standard files ENCFF497HEA and ENCFF246ZQE and K562 extended files ENCFF269DKY and ENCFF950FTI (https://www.encodeproject.org/). ABC predictions were downloaded from the Engreitz Lab AvgHiC ABC0.015 minus150 ForABCPaperV3 release (ftp://ftp.broadinstitute.org/outgoing/lincRNA/ABC/AllPredictions.AvgHiC.ABC0.015. minus150.ForABCPaperV3.txt.gz) and converted from hg19 to hg38 using the hg19-to-hg38 mapping invoked by ‘pyliftover.LiftOver("hg19", "hg38")‘, corresponding to the UCSC hg19ToHg38 chain resource (http://hgdownload.soe.ucsc.edu/goldenPath/hg19/liftOver/hg19ToHg38.over.chain.g z). Known motif sequences and cluster annotations were downloaded from JASPAR (https://jaspar.elixir.no/). SCENIC/cisTarget reference data were downloaded from the Aerts Lab resources: the human TF list (https://resources.aertslab.org/cistarget/tf_lists/allTFs_hg38.txt), the hg38 v10 clustered rankings database (https://resources.aertslab.org/cistarget/databases/homo_sapiens/hg38/refseq_r80/ mc_v10_clust/gene_based/hg38_10kbp_up_10kbp_down_full_tx_v10_clust.genes_v s_motifs.rankings.feather), and the motif-to-TF annotation table (https://resources.aertslab.org/cistarget/motif2tf/motifs-v10nr_clust-nr.hgnc-m0.001-o0.0.tbl). Compound annotations for Fig. 5 were curated from PubChem using PUG-REST and PUG-View (https://pubchem.ncbi.nlm.nih.gov/); the PubChem screening script and cached evidence tables are provided with the analysis code.

## Code availability

The analysis code is available at https://github.com/rikenbit/GenPerturb.

## Data availability

The perturbed scRNA-seq data were obtained from multiple sources: Norman et al. and Replogle et al. from scPerturb (http://projects.sanderlab.org/scperturb/); Srivatsan et al. from the sc-pert repository (https://github.com/theislab/sc-pert); Jiang et al. from GSE217460 (https://data.caltech.edu/records/2cjss-wgh69); and Martin et al. from GEO (GSE274113). Martin et al. Supplementary Tables S2 and S3 associated with the original article [12] were used for TF-sensitive ACR and TF-sensitive gene annotations. The GRCh38.p14 reference genome and GENCODE v46 basic gene annotation were downloaded from GENCODE release 46 (https://www.gencodegenes.org/human/release_46.html), including ‘GRCh38.p14.genome.fa.gz‘, ‘gencode.v46.chr_patch_hapl_scaff.basic.annotation.gff3.gz‘, and ‘gencode.v46.chr_patch_hapl_scaff.basic.annotation.gtf.gz‘; representative TSS coordinates were parsed from these annotation files by prioritizing Ensembl-canonical transcripts and using deterministic fallbacks when needed. Pretrained model weights were obtained through their model-specific download interfaces, including the official AlphaGenome JAX weights and Hugging Face repositories used by the PyTorch implementations. ENCODE-rE2G enhancer-gene predictions were downloaded from the ENCODE Portal, including the K562 standard files ENCFF497HEA and ENCFF246ZQE and K562 extended files ENCFF269DKY and ENCFF950FTI (https://www.encodeproject.org/). ABC predictions were downloaded from the Engreitz Lab AvgHiC ABC0.015 minus150 ForABCPaperV3 release (ftp://ftp.broadinstitute.org/outgoing/lincRNA/ABC/AllPredictions.AvgHiC.ABC0.015. minus150.ForABCPaperV3.txt.gz) and converted from hg19 to hg38 using the hg19-to-hg38 mapping invoked by ‘pyliftover.LiftOver("hg19", "hg38")‘, corresponding to the UCSC hg19ToHg38 chain resource (http://hgdownload.soe.ucsc.edu/goldenPath/hg19/liftOver/hg19ToHg38.over.chain.g z). Known motif sequences and cluster annotations were downloaded from JASPAR (https://jaspar.elixir.no/). SCENIC/cisTarget reference data were downloaded from the Aerts Lab resources: the human TF list (https://resources.aertslab.org/cistarget/tf_lists/allTFs_hg38.txt), the hg38 v10 clustered rankings database (https://resources.aertslab.org/cistarget/databases/homo_sapiens/hg38/refseq_r80/ mc_v10_clust/gene_based/hg38_10kbp_up_10kbp_down_full_tx_v10_clust.genes_v s_motifs.rankings.feather), and the motif-to-TF annotation table (https://resources.aertslab.org/cistarget/motif2tf/motifs-v10nr_clust-nr.hgnc-m0.001-o0.0.tbl). Compound annotations for Fig. 5 were curated from PubChem using PUG-REST and PUG-View (https://pubchem.ncbi.nlm.nih.gov/); the PubChem screening script and cached evidence tables are provided with the analysis code.

## Code availability

The analysis code is available at https://github.com/rikenbit/GenPerturb.

## Acknowledgements

We thank the members of the Omics AI Research Team, particularly Akihiro Matsushima, for their management of the IT infrastructure.

We thank the members of the Omics AI Research Team for their discussion of single-cell RNA-sequencing technologies.

## Funding

This work was partially supported by the Japan Science and Technology Agency, CREST Grant Numbers JPMJCR16G3, JPMJCR21N6, and JPMJCR1926, Japan.

This work was partially supported by the Japan Agency for Medical Research and Development (AMED) under Grant Numbers JP20bm0404024 and JP21bm0404073.

This work was partially supported by the RIKEN TRIP initiative (RIKEN TRIP-AGIS) in RIKEN and the Medical Research Center Initiative for High Depth Omics in Science Tokyo.

## Contributions

T.S. and I.N. designed and configured the various approaches used in this study.

T.S. performed the training of transfer learning. T.S. analyzed the data. T.S.. and I.N. prepared the figures and wrote the manuscript. All authors read and approved the final manuscript.

## Corresponding author

Correspondence to: Itoshi Nikaido

## Ethics declarations

### Ethics approval and consent to participate

Not applicable.

### Consent for publication

Not applicable.

### Competing interests

The authors report no competing interests.

## Notes

### Competing Interest Statement

The authors have declared no competing interest.

