## Supplemental table and figure for "GenPerturb: sequence-grounded interpretation of perturbation transcriptomes using pretrained genomic models"

### Supplementary information

### Supplementary Table 1. Comparative Overview of Computational Methods for Perturbation Response and Regulatory Interpretation

| Category | Method examples | Primary input | Primary output | Perturbation-aware | Quantitative expression output | Generalization to unseen perturbations | Sequence-level interpretability | Element–gene linking | Upstream TF |
|---|---|---|---|---|---|---|---|---|---|
| Sequence-based perturbation interpretation (this study) | GenPerturb | DNA sequence + target RNA-only Perturb-seq | Fitted expression outputs + differential attribution | ○ | ○ | × | ○ | ○ | ○ |
| Sequence-to-expression (bulk) | Enformer, Borzoi, AlphaGenome | DNA sequence | Expression / epigenome | × | ○ | × | ○ | ○ | ○ |
| Sequence-to-expression (single-cell) | GET, scooby, , Decima | DNA sequence + single-cell ATAC and/or RNA | Cell-type- / state-specific expression / epigenome | × | ○ | × | ○ | ○ | ○ |
| Perturb-seq analysis | MIMOSCA, scMAGeCK-LR | scRNA + gRNA | DEG / effect sizes | ○ | ○ | × | × | × | △ |
| gLM | HyenaDNA, Evo2, Nucleotide Transformer | DNA sequence | DNA functional annotation after fine-tuning | × | × | × | ○ | × | × |
| Base-resolution regulatory profile models | BPNet, ChromBPNet | DNA sequence + ATAC-seq or ChIP- | TF binding / chromatin accessi | × | × | × | ○ | × | ○ |

| | | | | | | | | | |
|---|---|---|---|---|---|---|---|---|---|
| | | seq profiles/peaks | bility profiles + attribution | | | | | | |
| ATAC-seq motif activity | chromVAR, ArchR, Signac | scATAC / peaks | Motif deviation / TF activity | △ | × | × | △ | × | ○ |
| Regulon / TF-activity inference | SCENIC, SCENIC+, RothEA, VIPER | scRNA (+ scATAC) | Regulons / TF activity scores | △ | △ | × | △ | △ | ○ |
| Enhancer–gene linking | ABC, rE2G | BED + cell-type-specific epigenomic annotations | Enhancer–gene scores | △ | × | × | × | ○ | × |
| Motif analysis | TOMTOM, MEME, gimme motifs | BED (genomic regions) | Motif discovery / matching | △ | × | × | ○ | × | ○ |
| Perturbation response prediction | GEARS, scGPT, CPA | Gene embeddings + perturbation labels | Expression prediction | ○ | ○ | ○ | × | × | × |
| Sequence-conditioned perturbation prediction | STRAND | DNA sequence (around perturbed locus) + scRNA (control cells) | Single-cell expression distribution after perturbation | ○ | ○ | ○ | × | × | × |

Note : Comparison with Related Methods

Supplementary Table 1 summarizes major computational methods for analyzing, modeling, and interpreting transcriptional responses to genetic perturbations, across multiple aspects including input, output, perturbation awareness, quantitative expression output, generalization to unseen perturbations, sequence-level interpretability, element–gene linking, and upstream transcription factor (TF) prioritization.

- ○: Provided as a native or primary function.
- △: Available only partially, indirectly, or through common downstream combination with additional resources.
- ×: Not provided by the method for the relevant comparison axis.

Below, we describe the roles and limitations of each category in the same order as Supplementary Table 1.

1. Sequence-based perturbation interpretation (GenPerturb)

GenPerturb prioritizes sequence-grounded regulatory hypotheses from observed RNA-only Perturb-seq responses. It links gene-level perturbation responses to candidate cis-regulatory elements, motif instances, and upstream regulatory programs on the same gene locus.

2. Sequence-to-expression models (bulk: Enformer, Borzoi, AlphaGenome) [1-3]

These models predict bulk gene expression and epigenomic signals from DNA sequence, integrating long-range cis context. Attribution analysis enables quantification of contributions from individual sequence elements and can support enhancer–gene linking. However, these attributions primarily reflect steady-state regulatory landscapes and do not directly capture perturbation-induced regulatory changes.

3. Sequence-to-expression models (single-cell: GET, scooby, Decima) [4-6]

GET integrates chromatin accessibility and sequence information to predict gene expression across many human cell types, while scooby extends pretrained sequence representations with cell-specific decoders to predict single-cell RNA and ATAC signals. Decima predicts cell-type- and state-resolved expression directly from sequence (RNA-only, after Borzoi-based training on large single-cell atlases) and can interpret cis-regulatory differences between observational states such as healthy versus disease. These models resolve cell-type- and state-specific regulation, but they condition expression on observational cell-type or disease covariates rather than fitting the transcriptional response to a defined genetic or chemical perturbation. Consequently, they do not produce the control-subtracted differential attribution that GenPerturb uses to prioritize perturbation-induced cis-regulatory changes on the same gene locus.

4. Perturb-seq analysis methods (MIMOSCA, scMAGeCK-LR) [7,8]

These methods estimate perturbation effect sizes via regression models using gRNA assignments while controlling for covariates. They are effective for statistical detection and quantification of perturbation effects but attribute expression changes directly to perturbation labels, without modeling upstream cis-regulatory mechanisms.

5. Genomic language models (gLMs: HyenaDNA, Evo2, Nucleotide Transformer) [9-11]

These models learn general-purpose sequence representations through self-supervised training on large DNA corpora. While adaptable to diverse downstream tasks via fine-tuning, they do not directly predict expression or perturbation responses and require task-specific training for regulatory interpretation.

6. Base-resolution regulatory profile models (BPNet, ChromBPNet) [12,13]

BPNet and ChromBPNet are models that learn base-resolution regulatory profiles from DNA sequences using experimental data such as ChIP-seq (for transcription factor binding) or ATAC-seq (for chromatin accessibility). They can also be used to identify important sequence features, such as regulatory motifs. However, these models predict assay-specific signals rather than gene expression changes, and they require corresponding ChIP-seq or ATAC-seq data for training or application.

7. ATAC-seq motif activity estimation (chromVAR, ArchR, Signac) [14-16]

chromVAR estimates transcription factor activity based on motif enrichment in accessible chromatin regions. Integrated frameworks such as ArchR and Signac extend this to multiomic analyses. While powerful, they require ATAC-seq data and depend on predefined peaks and known motif databases, rather than deriving regulatory contributions directly from sequence alone.

8. Regulon and TF-activity inference (SCENIC, SCENIC+, DoRothEA, VIPER) [17-20]

SCENIC infers regulons based on co-expression and motif enrichment, but it cannot directly connect transcription factors to specific cis-regulatory elements. SCENIC+ partially addresses this by incorporating chromatin accessibility information, but this requires additional ATAC-seq data. DoRothEA and VIPER estimate TF activity from gene expression using curated or inferred regulons, but they depend on predefined regulon resources and do not link activity to specific cis-regulatory sequence elements.

9. Enhancer–gene linking (ABC, rE2G) [21,22]

These methods estimate enhancer–gene interactions using chromatin contact and epigenomic data. While effective for mapping regulatory relationships in steady-state conditions, they rely on predefined regions and annotations and are not designed to detect perturbation-induced regulatory changes from RNA-only data. Public resource coverage is also cell-type dependent: the ABC prediction resource used in this study is distributed for 131 cell types and tissues, whereas rE2G extended genome-wide prediction resources are available for two cell lines, K562 and GM12878. Standard ENCODE-rE2G can be applied to additional biosamples with bulk ATAC-seq or DNase-seq, and optionally H3K27ac and Hi-C, but this still requires cell-type-matched epigenomic input or processing. Therefore, ABC and rE2G scores are valuable when appropriate annotations exist for the relevant cell type or a close proxy, but they are not uniformly available for arbitrary RNA-only Perturb-seq cell states.

10. Motif analysis (TOMTOM, MEME, GimmeMotifs) [23,24]

These methods perform motif discovery or matching within given genomic regions. Although perturbation-specific motif changes can be assessed by comparing region sets, results depend heavily on region selection. They do not directly provide element–gene linking or quantitative expression prediction.

11. Perturbation response prediction models (GEARS, scGPT, CPA) [25-27]

These models perform conditional prediction of gene expression in response to perturbations in a learned embedding space. By generalizing relationships between genes and perturbations, they can predict responses to unseen or combinatorial perturbations. However, since predictions are confined to expression space, they lack a mechanism to attribute responses to specific sequence elements such as motifs or regulatory regions.

12. Sequence-conditioned perturbation prediction models (STRAND) [28]
STRAND uses DNA sequence information as a feature to represent perturbations, enabling generalization to unseen perturbations with similar sequence context. However, this sequence information is not used to predict gene expression for each target gene. As a result, it cannot link gene expression changes to specific cis-regulatory elements that drive those responses.

Positioning of GenPerturb
GenPerturb provides a complementary interpretation layer between perturbation response prediction models, sequence-to-expression models, and regulatory inference methods. Whereas unseen-perturbation prediction models emphasize generalization in expression space, GenPerturb focuses on interpreting observed perturbation responses at the level of motifs and candidate cis-regulatory elements. This enables sequence-grounded regulatory hypothesis prioritization from RNA-only Perturb-seq datasets without requiring matched chromatin or multiome measurements in the target experiment.

Note on scope. Supplementary Table 1 maps where GenPerturb sits among related method families, organized by input, output, perturbation awareness, and interpretation target rather than by performance on a shared task. The listed methods differ in their required inputs—for example curated regulons (DoRothEA, VIPER), matched chromatin (SCENIC+, ABC, rE2G), or gene embeddings (GEARS, scGPT, CPA)—and in their primary outputs, so they are not all addressed to the same question on the same data. The comparison here is therefore positional rather than quantitative, with the columns intended to convey differences in scope rather than a ranking among methods.

**Supplementary Table 2. Dataset metadata used for training.**

| Perturbation | Cell type | Paper | Num. genes | Num. perturbations |
|---|---|---|---|---|
| CRISPRa | K562 | Norman et al. Science 2019 [26] | 12,105 | 220 |
| CRISPRi (gwps) | K562 | Replogle et al. Cell 2022 [27] | 8,078 | 1,545 |
| CRISPRi (essential gene) | K562 | Replogle et al. Cell 2022 [27] | 8,386 | 638 |
| CRISPRi | RPE1 | Replogle et al. Cell 2022 [27] | 8,600 | 409 |
| Compound | Myeloid | Jiang et al. Cell 2026 [28] | 12,319 | 496 |
| Compound | CD8T | Jiang et al. Cell 2026 [28] | 13,502 | 620 |
| Compound | CD4T | Jiang et al. Cell 2026 [28] | 12,839 | 537 |
| Compound | B cell | Jiang et al. Cell 2026 [28] | 14,095 | 290 |
| Compound | A549 | Srivatsan et al. Science 2020 [29] | 27,927 | 703 |
| Compound | K562 | Srivatsan et al. Science 2020 [29] | 27,991 | 683 |
| Compound | MCF 7.00 | Srivatsan et al. Science 2020 [29] | 23,224 | 740 |

| | | | | |
|---|---|---|---|---|
| CRISPR KO | Erythroid | Martin et al. Science 2025 [30] | 11,971 | 13 |

Note: "Num. perturbations" counts the total number of fitted output conditions, including the non-targeting control. For example, the Martin et al. value of 13 corresponds to 12 perturbations plus the non-targeting control, consistent with the 12 perturbation conditions evaluated in the enhancer and motif benchmarks (Fig. 3; Results).

**Supplementary Table 3. Provenance table of datasets and verifiable elements**

| Dataset | Modality | Cell type | Perturbation type | Perturbation modality | Motif evaluation | Enhancer evaluation |
|---|---|---|---|---|---|---|
| Martin et al. (Perturb-multiome) | Multiome (RNA + ATAC) | Erythroid lineage cells | TF perturbations | CRISPR KO | Yes (motif recovery and in silico motif mutation) | Yes (AUPRC, distance-stratified) |
| Norman et al. (Perturb-seq) | Single-cell RNA-seq | K562 cells | TF / other perturbations | CRISPRa | Yes (motif recovery and in silico motif mutation) | No |
| PRJNA1128171 / Metzner et al. (Multiome Perturb-seq) | Multiome (RNA + ATAC) | RPE-1 cells | Chromatin regulators | CRISPRi | No (not used for TF evaluation) | No (prepared for enhancer evaluation, but excluded because too few perturbations yielded sufficient links between perturbation-induced expression changes and ATAC peaks) |
| GSE277747 / Yan et al. (MultiPerturb-seq) | Multiome (RNA + ATAC) | BT16 cells with NIH3T3 spike-in | Chromatin regulators | CRISPRi | No (not used for TF evaluation) | No (excluded because, after applying the same preprocessing workflow as for |

| | | | | | | |
|---|---|---|---|---|---|---|
| | | | | | | other datasets, including Mixscape-based filtering, only one perturbation satisfied the downstream eligibility criteria) |
| GSE288996 / Shevade, Yang et al. (CAT-ATAC) | Multiome (RNA + ATAC) | iPSC and K562 cells | Chromatin regulators | CRISPRi | No (not used for TF evaluation) | No (prepared for enhancer evaluation, but excluded because the public data do not provide the sgRNA-to-perturbation correspondence table required to assign perturbation identities) |

Note: The Martin et al. erythroid Perturb-multiome dataset [33] served as the primary dataset with verifiable regulatory elements, used for motif recovery, in silico motif mutation, and distance-stratified AUPRC enhancer evaluation, while the Norman et al. K562 Perturb-seq dataset [29] was used for motif recovery and in silico motif mutation. PRJNA1128171 [34], GSE277747 [35], and GSE288996 [36] were examined as candidate perturbation-based multiome datasets (RNA + ATAC + gRNA) for enhancer-oriented validation. We evaluated whether GenPerturb prioritizes enhancers linked to genes with perturbation-induced expression changes using area under the precision–recall curve (AUPRC). Positive examples were defined as gene–enhancer or gene–peak links where both the gene showed differential expression and the linked ATAC peak or enhancer showed differential accessibility under the same perturbation. This evaluation requires (i) per-cell perturbation assignments and (ii) sufficient numbers of positive links per perturbation, as AUPRC is unstable when positives are sparse.

PRJNA1128171 was processed successfully but yielded too few positive links. Although this dataset includes chromatin-remodeler or histone-related perturbations with substantial ATAC changes, such changes were rarely linked to differentially expressed genes. After requiring both differential expression and linked ATAC/enhancer changes, only a small number of perturbations retained enough positives, rendering AUPRC evaluation unreliable due to lack of perturbation-level replication.

For GSE277747, the main limitation emerged earlier in the preprocessing workflow. When processed using the same workflow as the other datasets, including Mixscape-based filtering, only one perturbation satisfied the downstream eligibility criteria. This may reflect the low count depth per cell in the processed data, limited perturbation-induced gene expression changes, or too few cells with clearly detectable perturbation effects after filtering. Because this left insufficient perturbation-level replication for model training and validation, we did not advance this dataset to the training or enhancer-evaluation stage.

For GSE288996, same-cell RNA and ATAC data were available, but per-cell sgRNA assignments were missing. Only aggregated sgRNA counts were provided, preventing identification of perturbations at the single-cell level and thus perturbation-specific linkage between ATAC and gene expression changes.

Overall, none of the datasets met the requirements for this evaluation framework. More broadly, publicly available same-cell multiome datasets combining perturbation, RNA, ATAC, and gRNA with sufficient quality and scale remain limited, especially those providing enough linked differential ATAC peaks/enhancers and gene expression changes per perturbation. This constrains robust quantitative evaluation.

These limitations highlight the importance of methods that infer cis-regulatory elements from RNA-seq alone. Compared to ATAC-seq and multiome data, RNA-seq is more widely available across perturbation studies, large-scale atlases, and clinical datasets. Therefore, approaches that infer latent regulatory elements and transcriptional networks from gene expression offer a practical strategy under current data constraints.

### Supplementary Figure

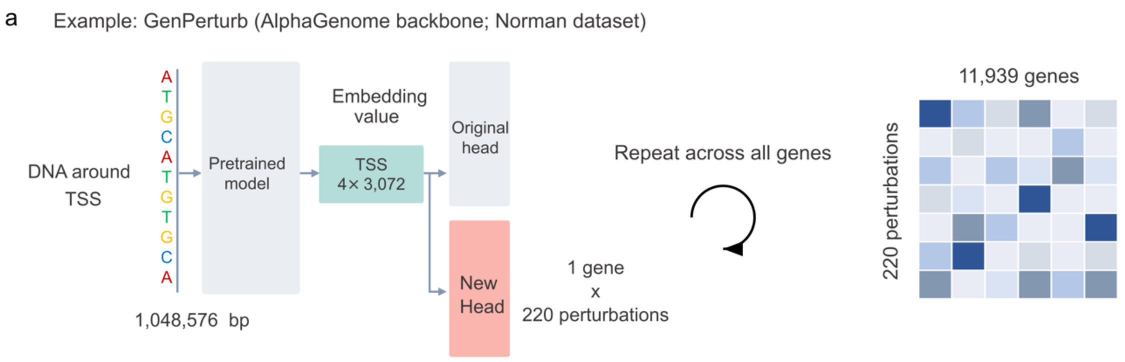


**Supplementary Figure S1. Sequence-embedding and prediction-head architecture used for GenPerturb transfer learning.**

Schematic of the GenPerturb architecture, shown here as an example using the Norman dataset and an AlphaGenome backbone; the same configuration is used for the other pretrained backbones and across transfer-learning strategies (feature extraction, LoRA, and full fine-tuning). For each gene, a 1,048,576-bp genomic sequence window centered on a representative transcription start site (TSS) is passed through the pretrained AlphaGenome backbone. The activations immediately preceding the original prediction head are retained for the four 128-bp bins surrounding the transcription start site, yielding a per-gene embedding of shape (4, 3,072), i.e. 12,288 dimensions. This embedding feeds two heads in parallel: the original pretrained prediction head, which reproduces the non-perturbed AlphaGenome output, and a newly fitted multi-output linear head followed by a ReLU activation, which is trained on the target Perturb-seq data. Each output channel of the new head corresponds to the non-targeting control or one observed perturbation condition in the target dataset and represents fitted per-gene log2(CPM + 1) expression. Applying the model gene by gene assembles a gene-by-perturbation expression matrix (11,939 genes across all data splits × 220 perturbation conditions in the depicted Norman example).

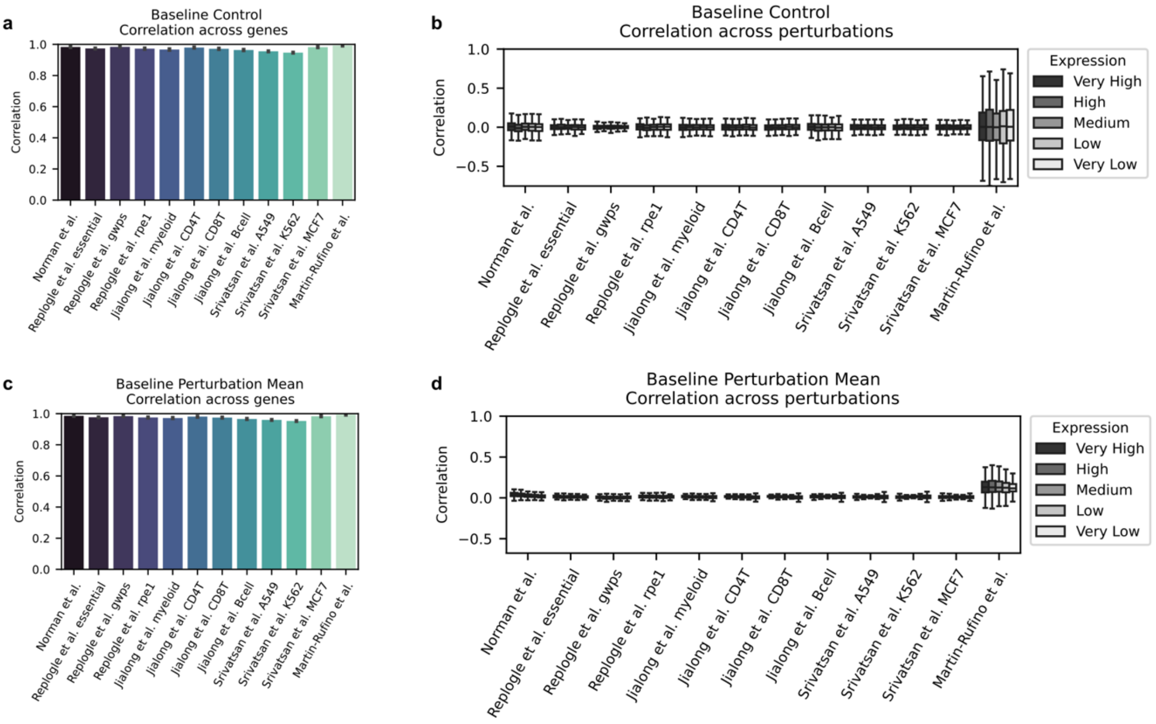


**Supplementary Figure S2. Simple expression baselines do not reproduce perturbation-dependent gene-wise variation.**

**a,** Across-gene agreement for the control-copy baseline, in which the non-perturbed expression profile is used in place of perturbation-specific predictions (Pearson r across genes, computed per perturbation).

**b,** Across-perturbation agreement for the control-copy baseline (Pearson r across perturbations, computed per gene; expression quintiles).

**c,** Across-gene agreement for the perturbation-mean baseline, in which the average expression profile across perturbations is used in place of perturbation-specific predictions (Pearson r across genes, computed per perturbation).

**d,** Across-perturbation agreement for the perturbation-mean baseline (Pearson r across perturbations, computed per gene; expression quintiles).

Although both baselines achieve high across-gene agreement, this performance largely reflects stable basal expression structure shared across perturbations. Neither baseline reproduces perturbation-dependent variation across genes, indicating that the across-perturbation signal captured by GenPerturb contains perturbation-specific transcriptional information beyond shared baseline or average response patterns. Across-perturbation variability was larger in the Martin et al. dataset because the analysis included only 12 perturbation conditions, resulting in less stable per-gene correlation estimates.

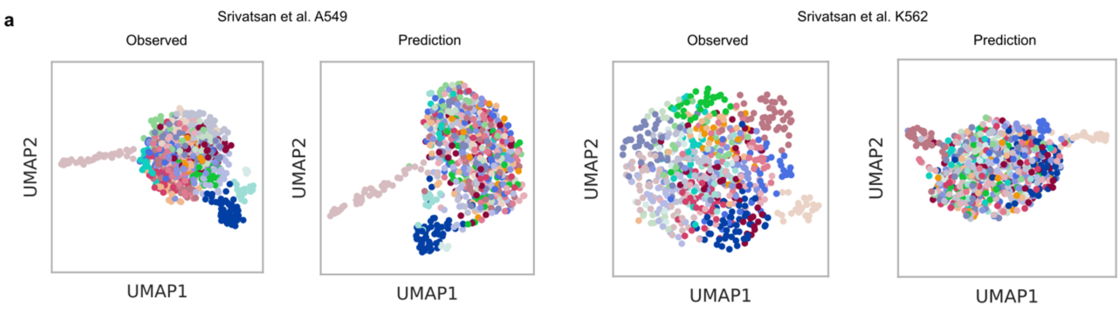


**Supplementary Figure S3. Low-dimensional organization of compound perturbation datasets with diffuse perturbation structure.**

a, Low-dimensional organization of observed and fitted expression profiles for compound perturbation datasets with more diffuse perturbation structure in the observed space (Srivatsan et al. A549 and K562). Dots in both observed and fitted

UMAPs are colored by Leiden clusters computed from the observed expression space, and the same observed-cluster colors are projected onto the fitted UMAPs. The fitted UMAPs preserve the weaker observed separation rather than imposing artificial cluster structure, complementing the datasets with pronounced perturbation structure shown in Fig. 2d.

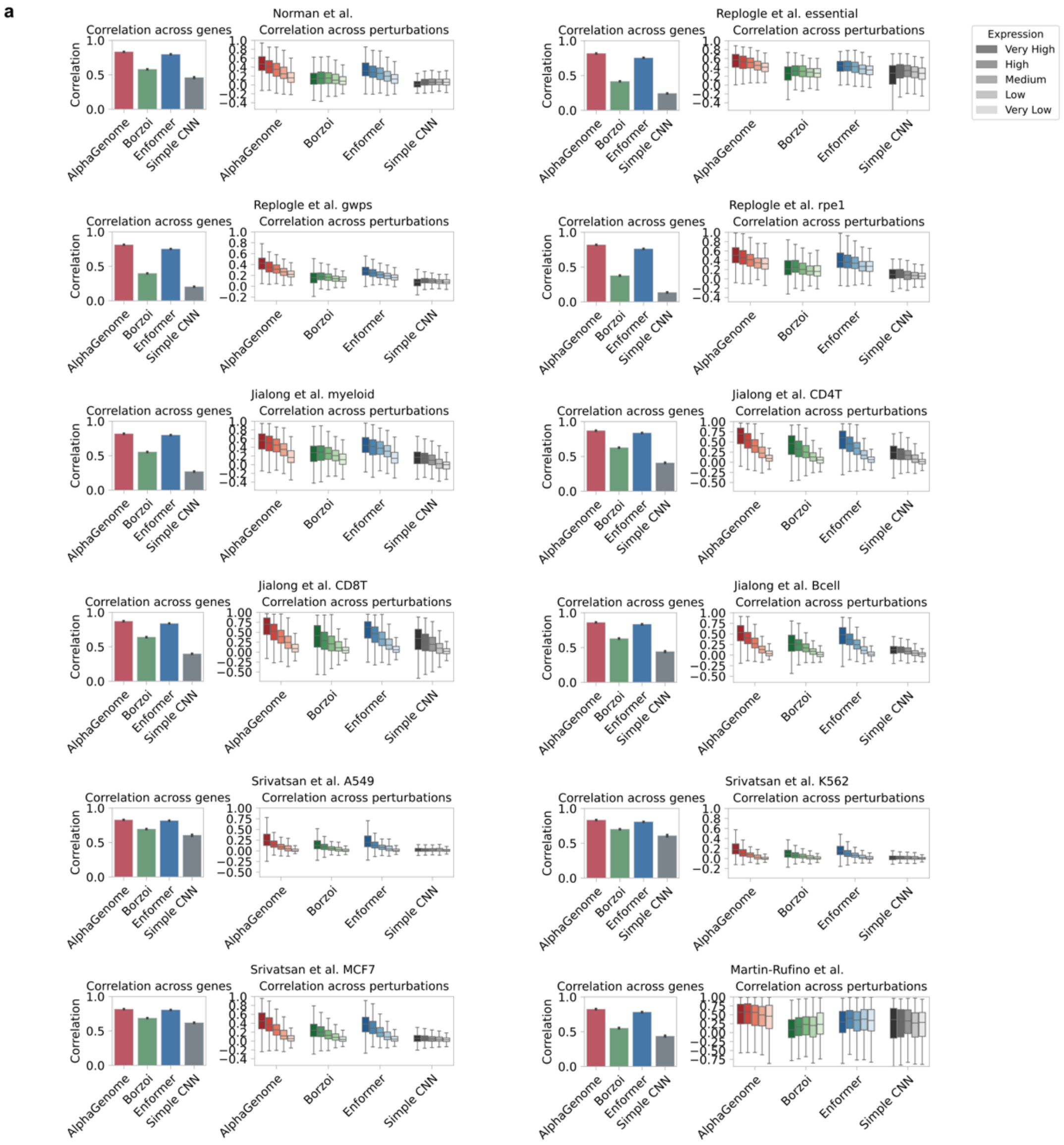


**Supplementary Figure S4. Comparison of pretrained sequence representation models and a non-pretrained CNN baseline for transfer learning.**

a, Across-dataset comparison of transfer learning using AlphaGenome, Borzoi, and Enformer sequence representations, together with a Simple CNN trained from scratch without large-scale regulatory pretraining (Pearson r; across-gene and across-perturbation evaluations; expression quintiles for across-perturbation panels; held-out test split). AlphaGenome-derived representations show stronger and more consistent agreement across datasets, supporting their use for capturing perturbation-dependent expression structure for downstream attribution analyses.

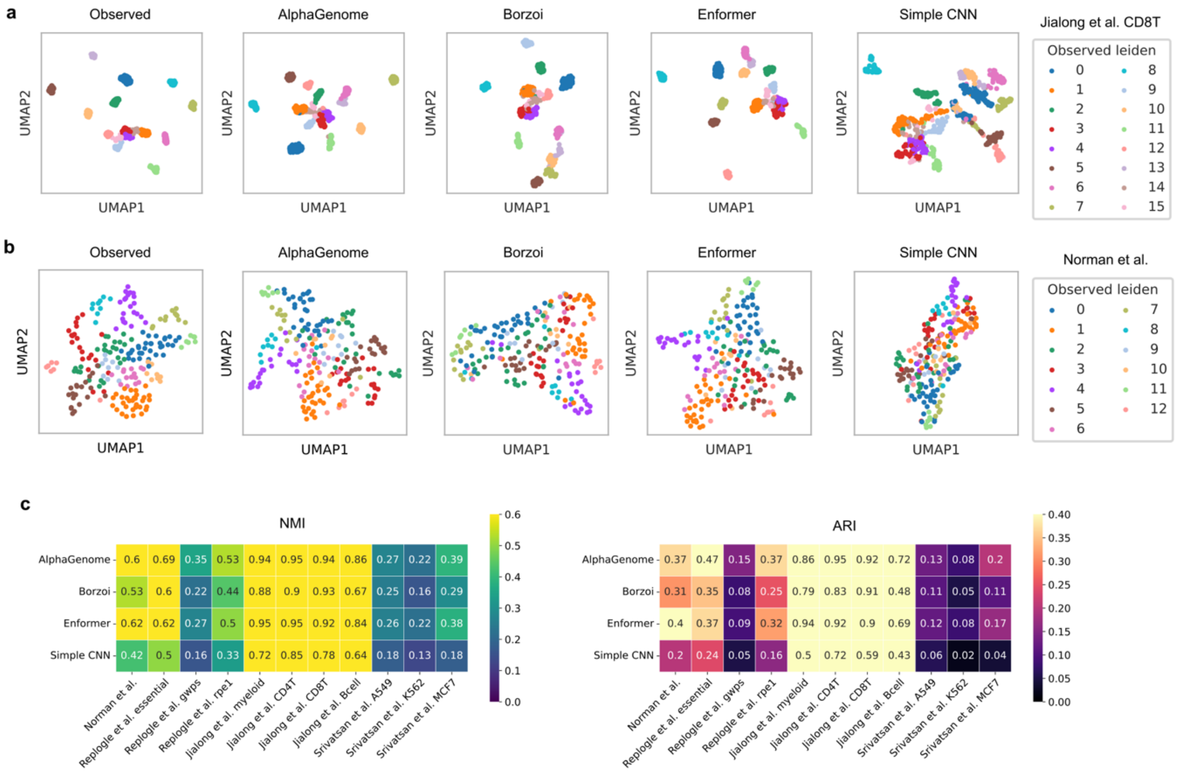


**Supplementary Figure S5. Comparison of sequence representation models in low-dimensional perturbation space.**

a, Low-dimensional organization of observed and fitted expression profiles for Jiang et al. CD8 T cells using transfer learning with AlphaGenome, Borzoi, Enformer, and a Simple CNN trained from scratch without large-scale regulatory pretraining. Observed Leiden cluster colors are projected onto the fitted UMAPs for each model (held-out test split).

b, Low-dimensional organization of observed and fitted expression profiles for Norman K562 perturbations using the same model set. Observed Leiden cluster colors are projected onto the fitted UMAPs for each model (held-out test split). c, Cluster concordance between observed and fitted expression spaces across models and datasets (ARI, NMI; observed vs fitted Leiden partitions; held-out test split). AlphaGenome-derived representations more effectively preserve perturbation-state geometry and fitted cluster structure than alternative pretrained representations or the non-pretrained Simple CNN baseline.

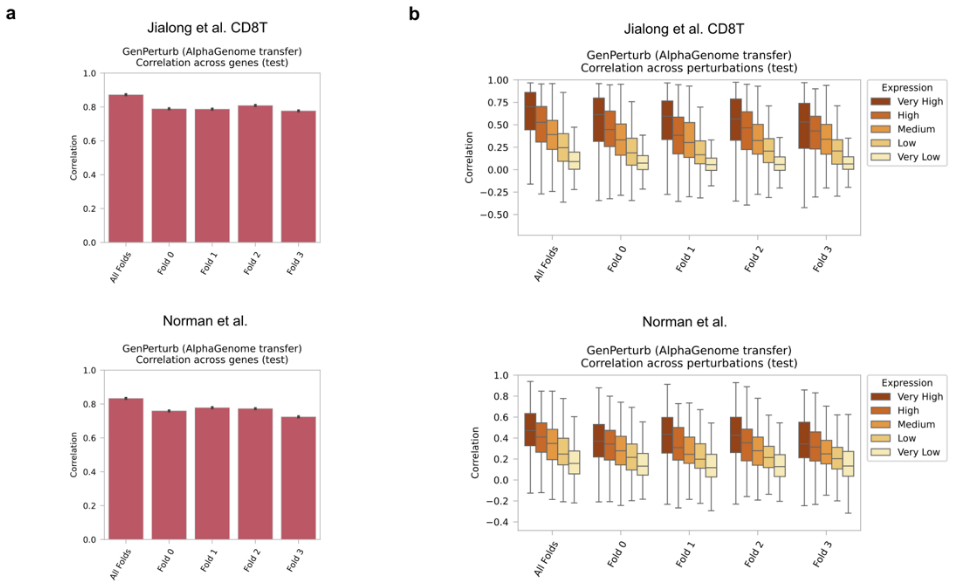


**Supplementary Figure S6. Stability of perturbation-dependent expression patterns across independent train/test splits.**

Multifold evaluation of AlphaGenome-based GenPerturb fitting using fold-specific pretrained AlphaGenome checkpoints and matched train/validation/test splits derived from the original AlphaGenome release. The all-fold comparator used the all-fold pretrained checkpoint and the same downstream prediction-head split as fold_0. a, Across-gene agreement across independent fold splits and the corresponding all-fold-checkpoint comparator (Pearson r across genes, computed per perturbation).

b, Across-perturbation agreement across independent fold splits and the corresponding all-fold-checkpoint comparator (Pearson r across perturbations, computed per gene; expression quintiles).

Results were consistent across fold-specific pretrained checkpoints and closely matched the all-fold-checkpoint comparator, supporting the stability of the perturbation-dependent expression structure retained by GenPerturb fitting.

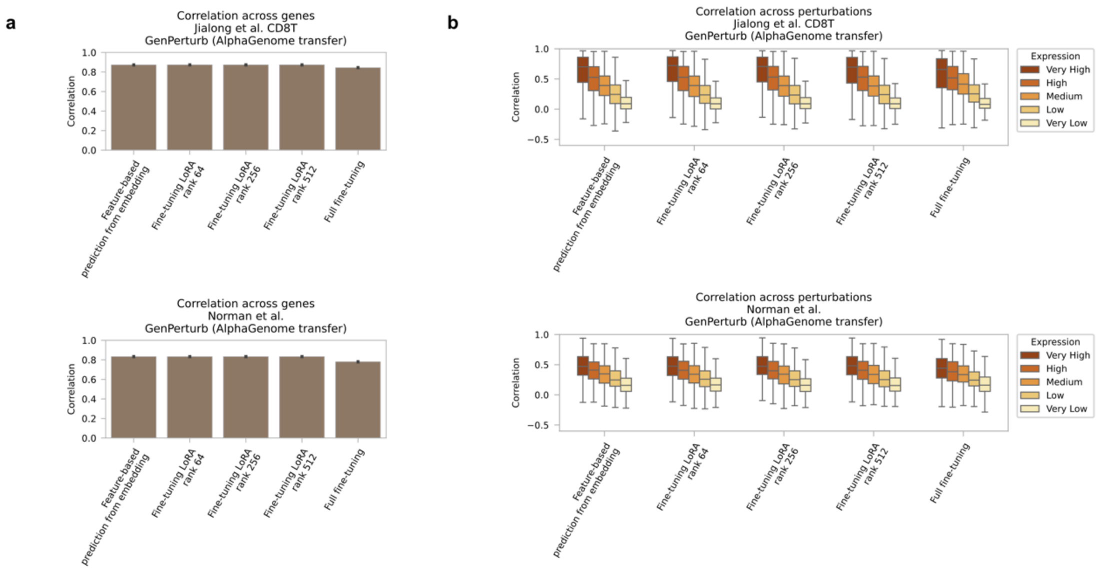


**Supplementary Figure S7. Comparison of transfer-learning strategies for pretrained sequence representations.**

Comparison of full fine-tuning, LoRA-based tuning, and feature extraction using AlphaGenome-derived representations.

a, Across-gene agreement for each fitting strategy (Pearson r across genes, computed per perturbation).

b, Across-perturbation agreement for each fitting strategy (Pearson r across perturbations, computed per gene; expression quintiles).

The transfer-learning strategies yielded broadly comparable agreement across datasets. Feature extraction was therefore used in subsequent analyses because it retained perturbation-dependent expression structure while substantially reducing computational cost. On a single NVIDIA H200 GPU, feature extraction required approximately one day for one-time precomputation of pretrained sequence embeddings followed by approximately 20 minutes for fitting the prediction head, whereas LoRA and full fine-tuning each required approximately one week per run.

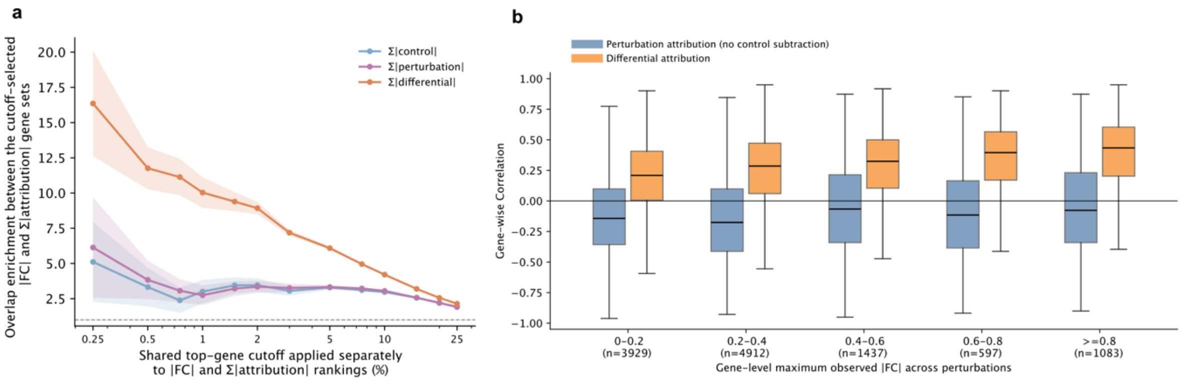


**Supplementary Figure S8. Gene-level attribution–fold-change concordance analysis supporting control-subtracted differential attribution.**

GenPerturb was fitted to the Martin et al. erythroid Perturb-multiome dataset, and gene-level attribution summaries were computed from genome-wide attribution profiles prior to enhancer benchmark evaluation.

a, Top-k overlap enrichment between genes ranked by observed absolute log fold change and genes ranked by summed absolute attribution magnitude. Rankings were computed separately for control-output attribution, perturbation-output attribution, and control-subtracted differential attribution within each perturbation, and enrichment was summarized across top-gene cutoffs. Lines show the mean across perturbations and shaded bands show the standard error.

b, Gene-wise Spearman correlation between observed absolute log fold change and attribution magnitude across perturbations, stratified by each gene's maximum observed absolute fold change. Perturbation-output attribution without control subtraction is compared with control-subtracted differential attribution. Box plots summarize the distribution across genes in each response-magnitude bin; sample sizes indicate the number of genes in each bin.

a

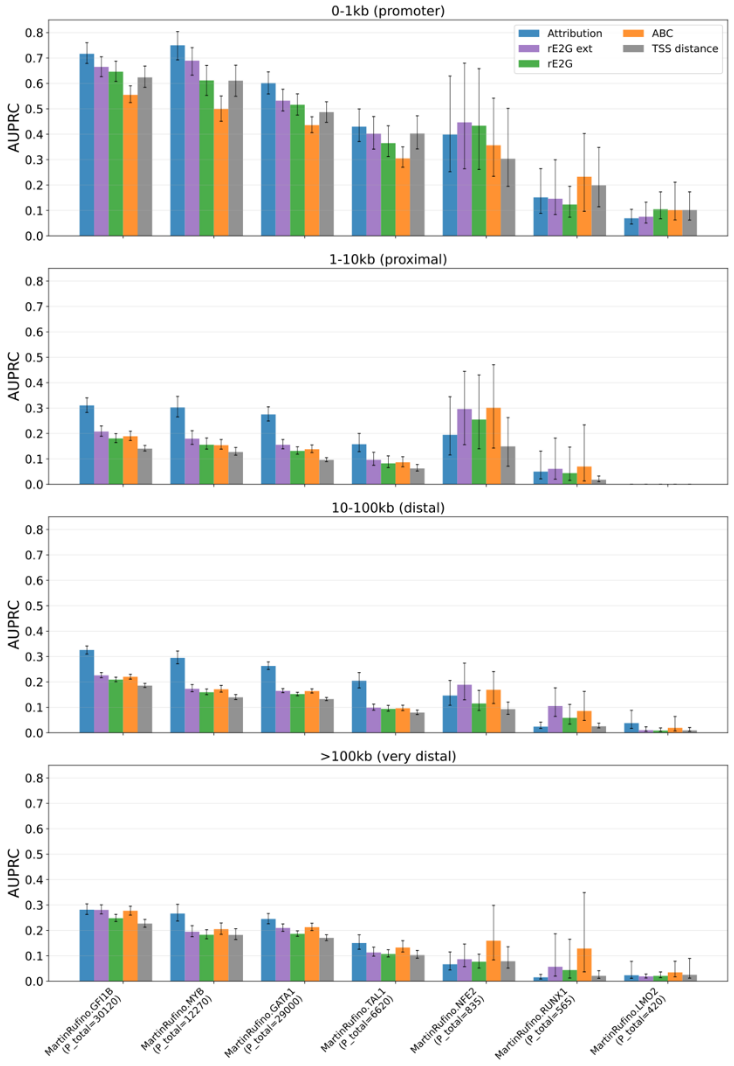


**Supplementary Figure S9. Per-perturbation enhancer prioritization across distance strata.**

a, Per-perturbation enhancer prioritization benchmark using perturbation-resolved TF-sensitive accessible chromatin regions (Martin et al. Table S2) as reference positives in the erythroid Perturb-multiome dataset (AUPRC; evaluated separately for each perturbation and distance stratum; |log2FC| ≥ 0.50 for gene-expression fold change; ≥10 positive peaks per distance bin). Differential attribution, rE2G extended, rE2G, ABC, and TSS-distance scores were compared across promoter-proximal (<1 kb), proximal (1–10 kb), distal (10–100 kb), and very distal (>100 kb) regulatory regions. Whiskers indicate within-perturbation 95% bootstrap confidence intervals obtained by resampling candidate peaks with replacement (1,000 iterations). Performance varied across perturbations and distance strata. Perturbations with larger numbers of positive enhancer candidates generally showed higher AUPRC and often favored differential attribution, whereas sparse-positive settings sometimes favored ABC or rE2G extended. These results suggest that model-derived attribution and annotation-based enhancer-linking methods provide complementary regulatory evidence rather than a single universally optimal ranking strategy.

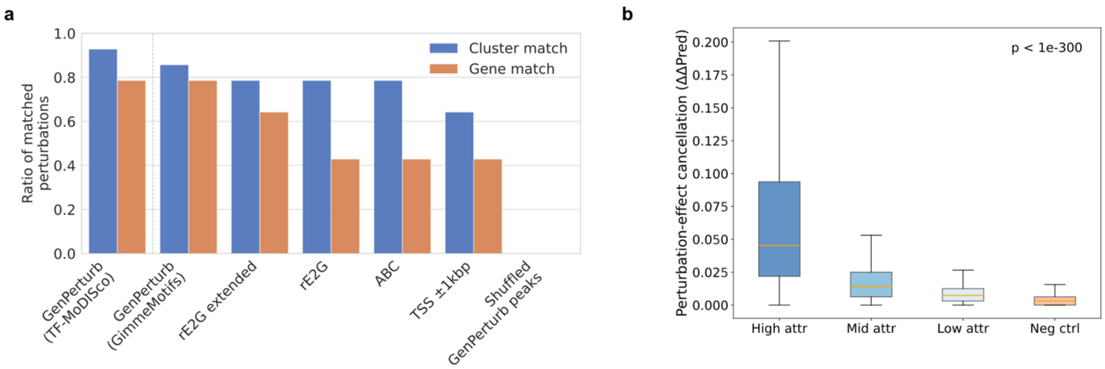


**Supplementary Figure S10. Motif recovery and model-internal mutation analysis in an independent CRISPRa dataset.**

GenPerturb was fitted to the Norman CRISPRa Perturb-seq dataset to test whether similar motif-level patterns could be reproduced across perturbation modalities and dataset contexts.

**a,** Motif recovery analysis comparing attribution-derived motifs and annotation-based regulatory regions (evaluated separately for each perturbation and motif-discovery method; TF-MoDISco and GimmeMotifs enrichments filtered at q ≤ 0.05; gene match or JASPAR transcription factor family cluster match to the perturbed transcription factor). GenPerturb attribution was evaluated using TF-MoDISco directly on differential attribution profiles and using GimmeMotifs known-motif enrichment on attribution peak regions, within the same motif-enrichment framework applied to annotation-based regions.

**b,** In silico mutation analysis of TF-MoDISco-identified motif instances whose Tomtom annotation matches the perturbed transcription factor or its JASPAR cluster ($q \leq 0.05$), stratified by differential attribution strength (5 mononucleotide-preserving random shuffle seeds averaged per mutated interval; groups = tertiles of attribution strength plus paired length-matched negative controls at motif-absent loci; one-sided Mann-Whitney U for high vs negative control where shown). High-attribution motif instances showed larger changes in fitted outputs upon sequence randomization than low-attribution or negative-control regions, supporting a consistent relationship between attribution magnitude and model-internal sequence sensitivity across perturbation contexts.

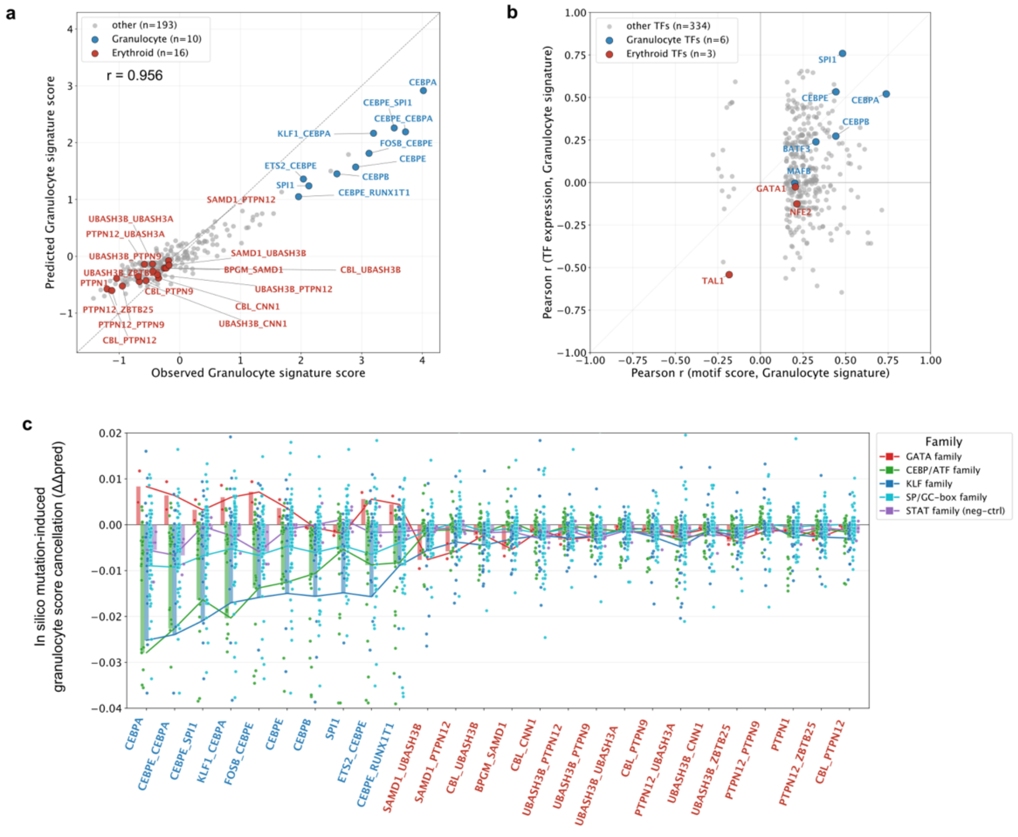


**Supplementary Figure S11. Granulocyte differentiation axis analysis in the Norman dataset.**

The lineage-axis analysis was extended from the erythroid axis to a granulocyte differentiation axis.

**a,** Comparison of observed and fitted granulocyte signature scores across perturbations using the granulocyte signature from the original Norman study (Pearson r). Each dot represents a perturbation; red dots indicate perturbations annotated as erythroid differentiation and blue dots indicate perturbations annotated as granulocyte differentiation in the original study.

**b,** Relationship between TF-MoDISco-derived motif significance scores (-log10 q) and TF mRNA expression along the granulocyte axis. For each motif or TF family, the y-axis shows the correlation between TF mRNA expression and the granulocyte signature score (Pearson r), and the x-axis shows the correlation between the motif significance score and the granulocyte signature score.

**c,** In silico mutation of granulocyte marker gene–associated and TF-MoDISco–identified motif instances (5 mononucleotide-preserving random shuffle seeds averaged per motif instance). Each dot represents a motif instance, and the y-axis shows the change in granulocyte marker-gene signature score after motif mutation (ΔΔpred). X-axis labels are colored using the original Norman study annotations, with erythroid differentiation perturbations in red and granulocyte differentiation perturbations in blue.

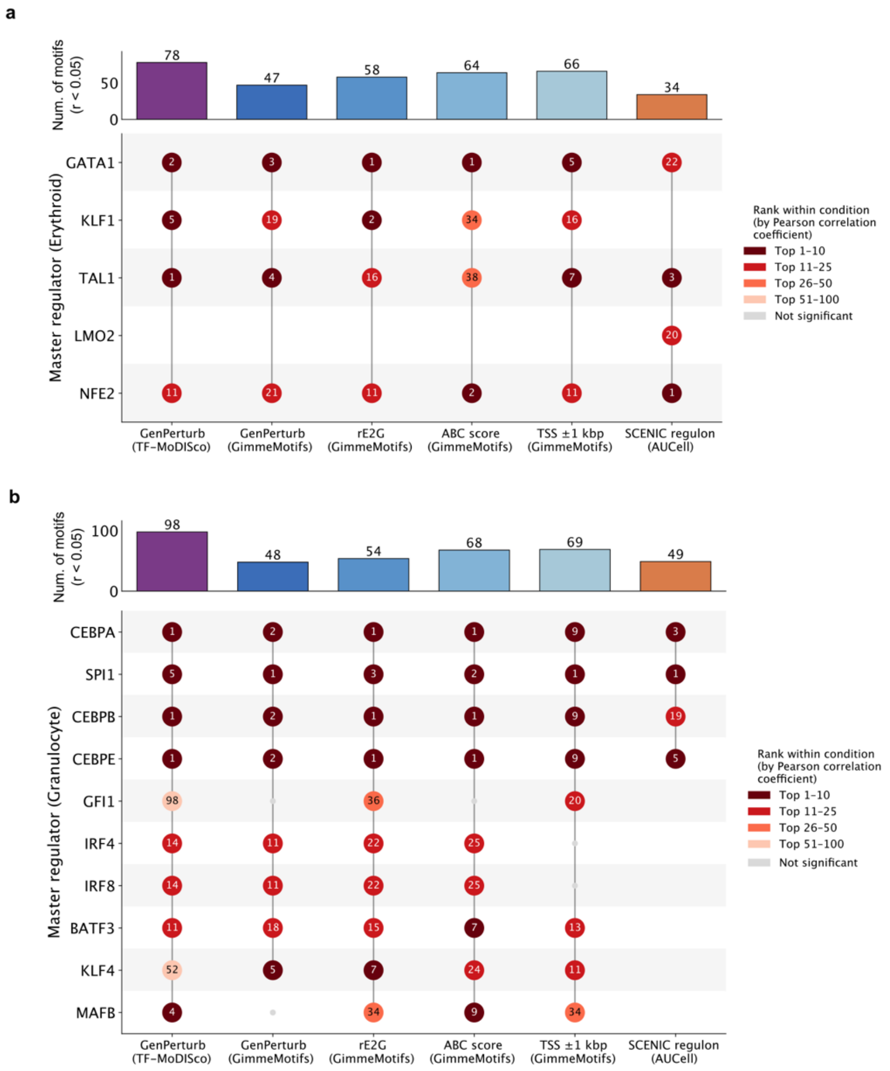

**Supplementary Figure S12. Comparison of lineage regulator ranking across attribution, enhancer-link, and regulon evidence.**

Comparison of candidate lineage regulators identified by GenPerturb attribution, motif enrichment over GenPerturb-derived regions, rE2G, ABC, TSS-proximal annotations, and SCENIC regulon activity. Regulator ranking was based on the absolute Pearson correlation coefficient between lineage signature scores and regulator-associated signals within each method. Numbers indicate regulator rank within each source. Top panels show the number of significant motif associations detected in each method (nominal $p < 0.05$).

**a,** Erythroid regulator ranking comparison. LMO2 was detected only by SCENIC because no corresponding LMO2 motif was included in the JASPAR database used for the motif-based analyses.

**b,** Granulocyte regulator ranking comparison.

Different methods showed partially overlapping regulator rankings, reflecting differences in attribution strategy, enhancer-link annotation, motif enrichment, and regulon inference. These approaches provide complementary evidence layers for interpreting lineage-associated regulatory programs.
